# Concept Recognition as a Machine Translation Problem

**DOI:** 10.1101/2020.12.03.410829

**Authors:** Mayla R Boguslav, Negacy D Hailu, Michael Bada, William A Baumgartner, Lawrence E Hunter

**Affiliations:** Computational Bioscience Program, University of Colorado Anschutz Medical Campus, 12635 East Montview Blvd, Aurora, CO 80045, USA

**Keywords:** Concept Recognition, Machine Translation, Named Entity Recognition, Named Entity Normalization, Computational Resources

## Abstract

**Background:** Automated assignment of specific ontology concepts to mentions in text is a critical task in biomedical natural language processing, and the subject of many open shared tasks. Although the current state of the art involves the use of neural network language models as a post-processing step, the very large number of ontology classes to be recognized and the limited amount of gold-standard training data has impeded the creation of end-to-end systems based entirely on machine learning. Recently, Hailu et al. recast the concept recognition problem as a type of machine translation and demonstrated that sequence-to-sequence machine learning models had the potential to outperform multi-class classification approaches. Here we systematically characterize the factors that contribute to the accuracy and efficiency of several approaches to sequence-to-sequence machine learning.

**Results:** We report on our extensive studies of alternative methods and hyperparameter selections. The results not only identify the best-performing systems and parameters across a wide variety of ontologies but also illuminate about the widely varying resource requirements and hyperparameter robustness of alternative approaches. Analysis of the strengths and weaknesses of such systems suggest promising avenues for future improvements as well as design choices that can increase computational efficiency with small costs in performance. Bidirectional Encoder Representations from Transformers for Biomedical Text Mining (BioBERT) for span detection (as previously found) along with the Open-source Toolkit for Neural Machine Translation (OpenNMT) for concept normalization achieve state-of-the-art performance for most ontologies in CRAFT Corpus. This approach uses substantially fewer computational resources, including hardware, memory, and time than several alternative approaches.

**Conclusions:** Machine translation is a promising avenue for fully machine-learning-based concept recognition that achieves state-of-the-art results on the CRAFT Corpus, evaluated via a direct comparison to previous results from the 2019 CRAFT Shared Task. Experiments illuminating the reasons for the surprisingly good performance of sequence-to-sequence methods targeting ontology identifiers suggest that further progress may be possible by mapping to alternative target concept representations. All code and models can be found at: https://github.com/UCDenver-ccp/Concept-Recognition-as-Translation.

## Background

Automated recognition of references to specific ontology concepts from mentions in text (hereafter “concept recognition”) is a critical task in biomedical natural language processing (NLP) and the subject of many open shared tasks, including BioCreAtIve [1], the BioNLP Open Shared Tasks (BioNLP-OST) [2], and the recent Covid-19 open research dataset challenge [3]. All of these shared tasks provide data, evaluation details, and a community of researchers, making them very useful frameworks for further development of such tasks. We chose to focus on the Concept Annotation Task of the CRAFT Shared Tasks at BioNLP-OST 2019 (CRAFT-ST) as our framework, not only because it is the most recent shared task involving concept recognition, but also because of the richness of data in CRAFT. All methods here can be applied to other corpora as well. The current state of the art (*e.g.*, [4]) involves the use of neural network language models as a post-processing step, as the very large number of ontology classes to be recognized and the limited amount of gold-standard training data has impeded the creation of end-to-end systems based entirely on machine learning. Furthermore, the goal of running such systems over the entirety of the vast biomedical literature (with more than one million new articles per year indexed in PubMed) means that the efficiency of such systems is important, as well as their accuracy [5].

Recently, Hailu et al. [6] recast the concept recognition problem as a type of machine translation and demonstrated that sequence-to-sequence machine learning models had the potential to outperform multi-class classification approaches. Here we systematically characterize the factors that contribute to the accuracy and efficiency of several approaches to sequence-to-sequence machine learning. The best-performing sequence-to-sequence systems perform comparably with the current state of the art (occasionally extending the state of the art by a modest degree), and some offer substantial efficiencies in the time and computational resources required for tuning and training. Furthermore, our analysis of the strengths and weaknesses of such systems suggests promising avenues for future improvements as well as design choices that can increase computational efficiency at a small cost in performance.

Concept recognition poses many difficult computational challenges. The target of most biomedical concept recognition efforts have been the Open Biomedical Ontologies, such as the Gene Ontology [7] or the Human Phenotype Ontology [8], each of which contain more than 10,000 specific concepts. Treating concept recognition as a classification task therefore requires tens of thousands of classes, rendering machine learning approaches impractical. Design principles for the Open Biomedical Ontologies require semantically coherent definitions, regardless of the variability in textual expression [9]. For example, the Gene Ontology Cellular Component class GO:0005886 might be textually referenced by “plasma membrane”, “cell membrane”, “cytoplasmic membrane”, or “plasmalemma”, among others, *e.g.*, abbreviations such as “PM”. All concept recognition methods must cope with challenges related to the variability and ambiguity of human language. Not only are there many lexical variants that refer to the same ontological concept, but each of those words have morphological variants (*e.g.*, nucleus, nuclei, nuclear, nuclearly). Furthermore, many individual words are ambiguous, depending on the surrounding context for the proper mapping to an ontological class; for example, the word “nucleus” can refer to an atomic nucleus (CHEBI:33252), a cell nucleus (GO:0005634), or an anatomical nucleus (UBERON:0000125), among other senses, depending on the surrounding context.

In the NLP literature, concept recognition is often divided into two tasks that are performed separately and differently (see Figure 1): Span detection (also referred to as named entity recognition or mention detection), which delimits a particular textual region that refers to some ontological concept (*i.e.*, a text mention); and concept normalization (also referred to as named entity normalization or entity linking), which identifies the specific ontological class to which the textual region or mention refers (an ontology class ID). It is possible to approach these problems jointly with a single system, but evaluation approaches in shared tasks generally score them separately and then combine the scores for the full system. Here we approach the problem separately employing very different methods for each task. Thus, going forward, we split every section into span detection and concept normalization, and both review them separately and together as the full system, aligning with the framework of the CRAFT-ST.

**Figure 1.**
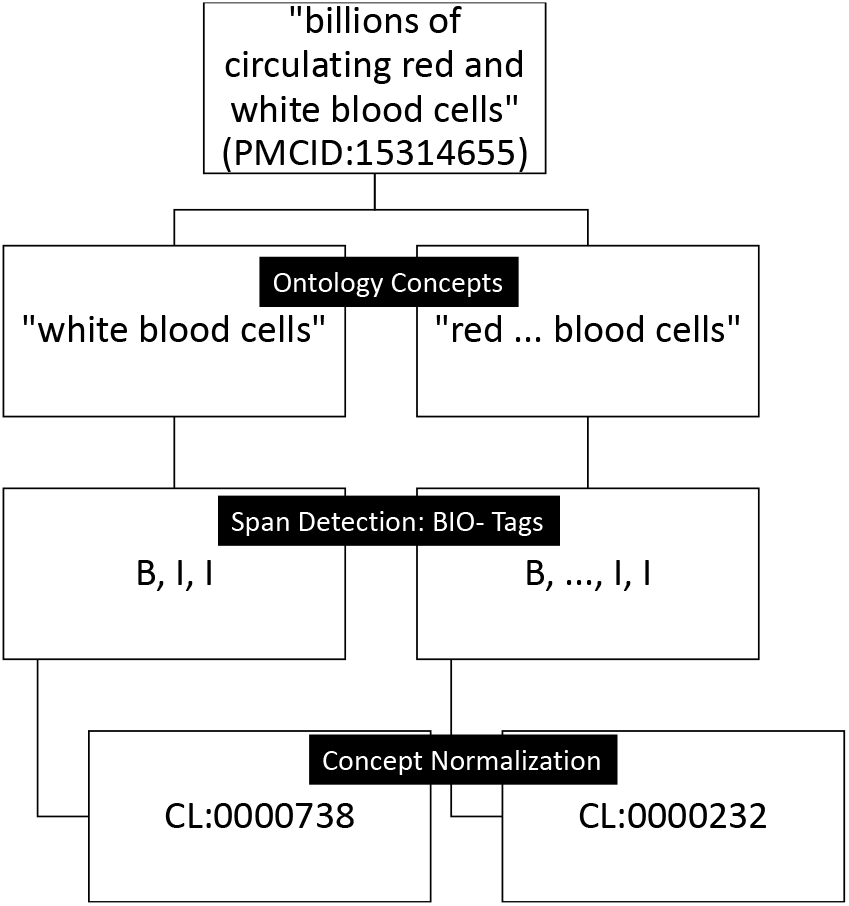
Example of the full translation pipeline. Each step is seen as a translation problem. The input is text and the final output is the ontology class identifiers for each detected text mention.

Span detection has long been conceived of as a sequence-to-sequence analysis task, with outputs defined as a sequence of tags identifying the beginning words or characters of mentions (labelled “B”), words or characters inside mentions (labelled “I”) and words or characters outside of mentions (labelled “O”), referred to as BIO tags [10]. We evaluated the most widely used and high-performing sequence-to-sequence algorithms for span detection to determine the best-performing algorithm as well as options for low-resource settings, specifically, conditional random fields (CRFs) [11] (which may be utilized with limited resources), bidirectional long short-term memory networks (BiLSTMs) [12, 13] (which require substantial resources, spurring us to evaluate whether hyperparameter settings tuned on a simple model can be reused in more complex models to save some resources), and language models (in particular, ELMo [14] and BioBERT [15],which are the current state of the art in NLP). In our evaluation, these systems show widely varying performances and resource requirements, and the performance is not strongly correlated with resource requirements. Furthermore, though they are relatively rare, we attempt to identify discontinuous concept mentions, *i.e.*, mentions that span two or more discontinuous regions of text. A few researchers have focused specifically on this problem recently [16, 17], while others have chosen to ignore them [4, 6]. We propose a simple extension to BIO tags that capture at least some discontinuous spans for almost all ontologies.

The novel contribution of Hailu et al. [6] was to use sequence-to-sequence methods for concept normalization, *i.e.*, to map detected spans to specific ontology class identifiers, both expressed as character sequences. Hailu et al. [6] investigated several alternative approaches and found that the popular Open Neural Machine Translation system (OpenNMT) [18] was the best-performing, although many other neural machine translation systems have very similar performance [19]. Thus, we continue the focus solely on OpenNMT for this exploration because the good performance of sequence-to-sequence approaches to normalization with the target being the ontology class identifier (*e.g.*, GO:0005634) is surprising, as the identifiers are intended to be arbitrary and without semantic content. We explore the reasons underlying this surprisingly good performance by evaluating a variety of alternative identifier schemes, suggesting that there are, in fact, semantic signatures in the class identifiers.

The performance of machine learning methods in NLP depends crucially on hyperparameter selection (as does the performance of many dictionary- and rule-based methods, see, *e.g.*, [20]). Large computational resource and time requirements, particularly for repeated training and testing under different hyperparameterizations, can limit the extent of hyperparameter search to find optimal values. Here we report on extensive studies of alternative methods and hyperparameter selections in an exhaustive evaluation framework that has been previously developed for open shared tasks [2]. These results not only identify the best-performing systems and parameters across a wide variety of ontologies but are illuminating with regard to the widely varying resource requirements and hyperparameter robustness of alternative approaches.

### Related Work

There have been many approaches to concept recognition, including dictionary-based or rule-based methods, classifier-based methods, and hybrids of these, using a variety of text representations ranging from bags of words to word and character embeddings [21, 22]. We evaluate alternative text representations using a modified sequence-to-sequence BIO tag representation [10] to represent text mentions, including overlapping and discontinuous mentions (see Figure 1 for an example). Previous research has mostly ignored these complex mentions due to their rarity and difficulty, and only recently have some researchers tried specifically to tackle them by extending existing sequence tagging frameworks (as we do here), or by looking at a given sentence as a whole to determine the relationships between entity mentions (see [16, 17] for an overview of previous work). The method proposed here is simpler than previous methods and still captures some complex spans.

Dictionary-based and rule-based methods dominate concept recognition approaches for the Open Biomedical Ontologies due to the enormous number of concepts to identify. Funk et al. [20] performed a systematic evaluation of some of these dictionary-lookup systems finding that ConceptMapper [23, 24] generally performed the best. They not only identified the highest-performing systems but also the best parameter settings (finding they were not the default settings), for each of the ontologies used in an earlier version of the CRAFT corpus, achieving F1 scores between 0.14 and 0.83. We thus use ConceptMapper, run on the updated version of CRAFT, as a baseline model in comparison to OpenNMT (see https://github.com/UCDenver-ccp/Concept-Recognition-as-Translation-ConceptMapper-Baseline for code). Rule-based post-processing techniques have also been proposed, including Boguslav et al. [25], which used the results of an error analysis to extend the ConceptMapper system from Funk et al. [20], thereby identifying post-processing techniques that improved precision with at most modest costs in recall.

Recent advances in machine learning have resulted in many hybrid systems that apply machine-learning-based post-processing to dictionary-based systems. For example, Campos et al. [26] proposed a hybrid system employing dictionary matching and a machine learning system for biomedical concept recognition. Groza et al. [27] approached the task as an information retrieval problem and explored case sensitivity and information gain. Basaldella et al. [28] proposed a hybrid system named OntoGene’s Entity Recognizer (OGER), which focused first on high recall through a dictionary-based entity recognizer, followed by a high-precision machine learning classifier (see [29] for an updated version of this system). Furthermore, the group who developed this system had the highest-performing method in the 2019 CRAFT Shared Task [4] (UZH@CRAFT-ST), combining an updated version of OGER with two neural approaches, thereby tackling concept recognition as a single task instead of two. As we are tackling the same task through the same framework, we use their results as a baseline for the full concept recognition system.

Many recent publications use sequence-to-sequence approaches for span detection but do not attempt normalization. For example, Huang et al. [30] proposed a model based on a BiLSTM combined with a CRF for BIO tagging and achieved better tagging accuracy for part-of-speech tagging, chunking, and span detection than with a CRF alone. Throwing in character and word embeddings, Lample et al. [31] used the same neural architecture as Huang et al. for span detection. Adding a CNN to the mix, Ma et al. [32] proposed an end-to-end sequence-tagging model based on a BiLSTM-CNN-CRF approach. Thinking beyond any specific language, Gillick et al. [33] created an LSTM-based model that reads text as bytes and outputs the span annotations. Since they focus on the bytes, their representations and models generalize across many languages, creating the first multilingual named entity recognition system.

In the biomedical domain, Habibi et al. [34] applied the BiLSTM-CRF proposed by Lample et al. [31] for span detection on a wide range of biomedical datasets and found that their model outperformed the state-of-the-art methods. The same architecture was used by Gridach [35] to identify spans of genes and proteins. Zhao et al. [36] proposed a multiple-label strategy to replace the CRF layer of a deep neural network for detecting spans of disease mentions. To identify spans of chemicals, Korvigo et al. [37] applied a CNN-RNN network, Luo et al. [38] proposed an attention-based BiLSTM-CRF and Corbett et al. [39] explored a BiLSTM and CRF separately as well as combined to create ChemListem and further added transfer learning. Similar to Luo et al, Unanue et al. [40] used a BiLSTM-CRF to identify spans of drug names and clinical concepts, while Lyu et al. [13] proposed a BiLSTM-RNN model to detect spans of a variety of biomedical concepts, including DNA, genes, proteins, cell lines, and cell types. Wang et al. [41] applied multitask learning with cross-sharing structure using a BiLSTM-CNN-CRF model, which includes a BiLSTM that learns shared features between ten datasets with gene, protein, and disease categories, and a private BiLSTM specific for each task, borrowing their base model from Ma et al. [32].

More recent advances in deep learning for a variety of different NLP tasks, including span detection, have been achieved with language models. BERT [42], the science-specific language model SciBERT [43], and the biomedicine-specific language model BioBERT [15], all require little fine-tuning to perform well for named entity recognition. The other main language model, ELMo [44], and its biomedical equivalent [14], also perform well on named entity recognition. Peng et al. [45] evaluated both BERT and ELMo on the Biomedical Language Understanding Evaluation (BLUE) benchmark and found that BERT, with extra biomedicine-specific documents, outperformed ELMo. Even though none of this research attempted concept normalization, the success of these methods show the value of deep learning and language models for span detection.

Deep learning methods have also been applied to concept normalization, although generally in hybrid settings. For example, Li et al. [46] generated concept normalization candidates using a rule-based system and then ranked them using a convolutional neural network that harnessed semantic information. Liu et al. [47] used an LSTM to represent and normalize disease names. Similarly, Tutubalina et al. [48] used recurrent neural networks, including LSTMs, to normalize medical concepts in social media posts.

Deep learning has also been applied to machine translation methods. To the best of our knowledge, this is the first attempt to use machine translation for concept normalization, and so there is no prior literature. However, machine translation usually aims to translate from one language to another (see, *e.g.*, [19,49]). Recently, neural machine translation has proven superior to previous methods. Successful approaches belong to the family of encoder-decoders, which encode source text into fixed-length vectors from which a decoder generates translations [19]. This is a sequence-to-sequence method that maps sequences of characters or tokens in one language into sequences of characters or tokens in another language. Bahdanau et al. [50] introduced a key innovation by adding an attention mechanism to the decoder to relieve the encoder of needing to encode all information from the source text to fixed-length vectors. With this approach, the information can be spread throughout the sequence of text, and the decoder can select the most useful parts to predict the next character or token. It is this approach that Hailu et al. exploited [6] and the approach we further explore here.

## Methods

Our goal is to explore the performance, efficiency, and underlying reasons for the surprisingly good performance of our machine learning approach toward concept recognition, using the 2019 CRAFT Shared Task framework from the BioNLPOST task [51, 52]. This framework provides all data and an evaluation pipeline facilitating direct comparison to the best-performing system in that evaluation [4]. Per the setup of the CRAFT Shared Task, 67 full-text documents were provided as training with 30 unseen documents held out as an external evaluation set. Since each ontology (*e.g.*, the ChEBI chemical ontology, the Uberon anatomical ontology) presents different challenges and may benefit from different methods, hyperparameters and/or training regimes, all of the ten Open Biomedical Ontologies (OBOs) used to conceptually annotate the CRAFT version used here, were trained, tuned, and evaluated independently for both span detection and concept normalization. Furthermore, training all ontologies separately solves both the problem of overlapping ontology annotations from different ontologies and the multi-classification problem between ontologies, and it simplifies the process of adding new ontologies. For all ontologies, the goal is to optimize F1 score, the harmonic mean between precision and recall, and it is the metric for comparison of all models. We also take into consideration the resources needed for all models.

### Materials and Evaluation Platform

The Colorado Richly Annotated Full-Text (CRAFT) corpus [53–55] was used to train, tune, and evaluate our models. CRAFT has had four major releases. The CRAFT Shared Task used version 3.1.3, which was released in July 2019 [56]; we thus used this release for all tasks and we will refer to it as CRAFT without the version number. Version 3.1.3 includes 67 full-text documents with 30 documents held back, whereas the most recent version includes the previously held-back 30 documents for a total of 97 [57]. These versions are a significant improvement from version 2.0 with regard to the concept annotations. Among other changes, relative to version 2.0, the concept annotations were updated using newer versions of the ontologies, annotations were created based on the classes of two additional ontologies (the Molecular Process Ontology and the Uberon anatomical ontology), and extension classes of the OBOs were created and used to annotate the articles, resulting in a substantial increase in annotation counts. (The extension classes were created by the CRAFT semantic annotation lead for the concept annotations but are based on proper OBO classes; they were created for various reasons, particularly for the purposes of semantic unification of similar classes from different ontologies, unification of multiple classes that were difficult to consistently differentiate for annotation, and creation of similar but corresponding classes that were either easier to use or better captured the ambiguity of the annotated text mentions.) The total corpus is a collection of 97 full-text articles with a focus (though not exclusively) on the laboratory mouse that has been extensively marked up with both gold-standard syntactic and semantic annotations. Among the syntactic annotations, segmented sentences, tokens, and part-of-speech tags were used to extract features for all algorithms tested. The semantic annotations to single or multi-word concepts rely on ten Open Biomedical Ontologies (OBOs):

1. Chemical Entities of Biological Interest (ChEBI): compositionally defined chemical entities (atoms, chemical substances, molecular entities, and chemical groups), subatomic particles, and role-defined chemical entities (*i.e.*, defined in terms of their use by humans, or by their biological and/or chemical behavior)
2. Cell Ontology (CL): cells (excluding types of cell line cells)
3. Gene Ontology Biological Processes (GO_BP): biological processes, including genetic, biochemical/molecular-biological, cellular and subcellular, organ- and organ-system level, organismal, and multiorganismal processes
4. Gene Ontology Cellular Components (GO_CC): cellular and extracellular components and regions; species-nonspecific macromolecular complexes
5. Gene Ontology Molecular Function (GO_MF): molecular functionalities possessed by genes or gene products, as well as the molecular bearers of these functionalities (though note that only five of the proper classes of the ontology were used for annotation, while the corresponding extension classes were instead used for the large majority of the classes of this ontology)
6. Molecular Process Ontology (MOP): chemical reactions and other molecular processes
7. NCBI Taxonomy (NCBITaxon): biological taxa and their corresponding organisms; taxon levels
8. Protein Ontology (PR): proteins, which are also used to annotate corresponding genes and transcripts
9. Sequence Ontology (SO): biomacromolecular entities, sequence features, and their associated attributes and processes
10. Uberon (UBERON): anatomical entities; multicellular organisms defined in terms of developmental and sexual characteristics

For each of the ten ontologies used in the CRAFT Corpus, there are two annotation sets: a core set and a core+extensions set. The core set consists solely of annotations made with proper classes of the given OBO, and the core+extensions set consists of annotations with proper OBO classes as well as classes created as extensions of the ontologies. The unique identifier for an OBO class is a class identifier (class ID) that consists of the ontology namespace, a colon, and a unique number identifier; for example, NCBITaxon:10088 is the unique identifier for *Mus*, the taxonomic genus of mice. An extremely small subset of classes of the OBOs used for CRAFT concept annotation instead have textual IDs, *e.g.*, NCBITaxon:species, the NCBI Taxonomy class representing the taxonomic rank of species. Between and within ontologies the length of the class IDs can vary: CL, GO, MOP, SO and UBERON classes have seven-digit IDs (*e.g.*, CL:0000014 (germ line stem cell), GO:0016020 (membrane), MOP:0000590 (dehydrogenation), SO:0000040 (genomic clone), UBERON:0001004 (respiratory system)), while PR classes have nine-digit IDS (*e.g.*, PR:000000035 (BMP receptor-type 1A)). CHEBI class IDs range from one (*e.g.*, CHEBI:7 ((+)-car-3-ene)) to six (*e.g.*, CHEBI:139358 (isotopically modified compound)) digits, and NCBITaxon entry IDs (other than those representing taxonomic ranks) have between one (*e.g.*, NCBITaxon:2 (Bacteria)) and seven (*e.g.*, NCBITaxon:1000032 (*Enterobacter* sp. P19-19)) digits. For the extension classes, which are identifiable by their namespace prefixes always ending in “_EXT”, the class IDs are even more varied. For four of the OBOs used, one or more parallel hierarchies of extension classes were programmatically created for either the entire OBO or for one or more subhierarchies of the OBO so as to create corresponding classes that are more abstract and/or more straightforward to use for concept annotation; the class ID for such an extension class is the same as the original OBO class on which it is based with the exception of “_EXT” appended to the namespace, *e.g.*, GO_EXT:0004872 (bearer of signaling receptor activity), based on the original GO:0004872 (signaling receptor activity). In addition to these programmatically created extension classes, there are many manually created extension classes, which have entirely textual IDs of human-readable labels with the namespaces of the ontologies of which they are extensions appended with “_EXT”, *e.g.*, CHEBI_EXT:calcium. In the case of a manually created extension class that is an extension of more than one ontology, the namespace is an underscore-delimited concatenation of the namespaces of the extended ontologies, *e.g.*, GO_MOP_EXT:glycosylation. Of relevance to our concept normalization work, many of the manually created extension classes have long textual IDs, *e.g.*, GO_UBERON_EXT:innervation entity or process. More information about the CRAFT Corpus can be found at [58], and statistics regarding the annotations and annotation classes of the training and evaluation data sets can be seen in Tables 1 and 2, respectively. Note that only a very small number of proper GO_MF classes were used for annotation and that the corresponding extension classes were instead used for the large majority of classes in the GO_MF core+extensions annotation set (GO_MF_EXT).

**Table 1.** Statistics for the concept annotations in the training (67-document) and evaluation (30-document) data sets for all ontologies. Note: avg stands for average.

| Ontology | # training set annotations | avg/median # training set annotations per article | # evaluation set annotations | avg/median # evaluation set annotations per article |
| --- | --- | --- | --- | --- |
| ChEBI | 4548 | 68/45 | 2200 | 73/45 |
| ChEBI_EXT | 11915 | 178/142 | 5248 | 175/142 |
| CL | 4043 | 60/32 | 1749 | 58/32 |
| CL_EXT | 6276 | 94/64 | 2872 | 96/64 |
| GO_BP | 9280 | 139/108 | 3681 | 123/108 |
| GO_BP_EXT | 13954 | 208/158 | 5847 | 195/158 |
| GO_CC | 4075 | 61/33 | 1184 | 39/33 |
| GO_CC_EXT | 8495 | 127/91 | 3217 | 107/91 |
| GO_MF | 375 | 6/2 | 94 | 3/2 |
| GO_MF_EXT | 4070 | 61/34 | 1822 | 61/34 |
| MOP | 240 | 4/2 | 101 | 3/2 |
| MOP_EXT | 386 | 6/2 | 111 | 4/2 |
| NCBITaxon | 7362 | 110/90 | 3101 | 103/90 |
| NCBITaxon_EXT | 7592 | 113/97 | 3219 | 107/97 |
| PR | 17038 | 254/198 | 6409 | 214/198 |
| PR_EXT | 19862 | 296/246 | 7932 | 264/246 |
| SO | 8797 | 131/118 | 3446 | 115/118 |
| SO_EXT | 24955 | 372/341 | 9136 | 305/341 |
| UBERON | 12269 | 183/130 | 6551 | 218/130 |
| UBERON_EXT | 14910 | 223/165 | 7416 | 247/165 |

**Table 2.** Statistics for the concept annotation classes used in the training (67-document) and evaluation (30-document) data sets and for those added for additional training data for concept normalization for all ontologies. Note: avg stands for average.

| Ontology | # training set<br>annotation<br>classes | avg/median # training set<br>annotation classes per<br>article | # classes added<br>to training set | # evaluation<br>set annotation<br>classes | avg/median # evaluation<br>set annotation classes per<br>article |
| --- | --- | --- | --- | --- | --- |
| ChEBI | 1463 | 22/18 | 58214 | 627 | 21/20 |
| ChEBI_EXT | 2852 | 43/38 | 58439 | 1167 | 39/39 |
| CL | 581 | 9/7 | 2163 | 253 | 8/9 |
| CL_EXT | 651 | 10/8 | 2168 | 286 | 10/10 |
| GO_BP | 1586 | 24/21 | 29213 | 682 | 23/23 |
| GO_BP_EXT | 2511 | 37/33 | 29301 | 1090 | 36/37 |
| GO_CC | 677 | 10/9 | 4052 | 212 | 7/6 |
| GO_CC_EXT | 896 | 13/12 | 4086 | 296 | 10/9 |
| GO_MF | 49 | 1/1 | 10951 | 19 | 1/1 |
| GO_MF_EXT | 738 | 11/11 | 10031 | 377 | 13/12 |
| MOP | 85 | 1/1 | 3574 | 32 | 1/1 |
| MOP_EXT | 108 | 2/1 | 3578 | 40 | 1/1 |
| NCBITaxon | 690 | 10/9 | 1175661 | 315 | 11/9 |
| NCBITaxon_EXT | 757 | 11/10 | 1175682 | 346 | 12/10 |
| PR | 1278 | 19/18 | 213371 | 466 | 16/16 |
| PR_EXT | 1534 | 23/22 | 213531 | 588 | 20/19 |
| SO | 1216 | 18/18 | 2256 | 544 | 18/19 |
| SO_EXT | 3172 | 47/47 | 2405 | 1409 | 47/48 |
| UBERON | 2048 | 31/24 | 14057 | 1040 | 35/31 |
| UBERON_EXT | 2409 | 36/29 | 14113 | 1217 | 41/38 |

Performance of all systems is measured using the CRAFT Shared Task evaluation platform [51, 52]. Briefly, for the concept annotation task, the evaluation platform makes use of the method proposed by Bossy et al. [59], which incorporates flexibility in matching both the boundaries (*i.e.*, the start and end of a concept mention in the text) of a predicted concept to the reference, and in the predicted ontology class identifiers. This flexibility allows the scoring metric to assign partial credit to inexact matches in two different ways. Partial credit is assigned for overlapping boundaries using a Jaccard index scheme over the characters in the matches, and partial credit for inexact ontology class ID matches is computed using a semantic similarity metric that makes use of the hierarchical structure of the ontologies. The final scores for a given predicted concept are based on a hybrid of both the boundary and ontology class ID match scores, and include precision, recall, F1 score, and an aggregate score called the slot error rate (see Bossy et al. [59] for the exact equations). In order to facilitate reproducibility and comparison of future systems to those that participated in the 2019 CRAFT Shared Task, the evaluation platform is made available as a versioned Docker container. Version 4.0.1 0.1.2 was used to evaluate the systems described here. All evaluation information can be found at the CRAFT GitHub site (specifically at https://github.com/UCDenver-ccp/craft-shared-tasks/wiki/Concept-Annotation-Task-Evaluation).

Computational time and resource availability limited some potential evaluations. Some computations were performed on a contemporary laptop, but many required the use of an NIH-funded shared supercomputing resource [60] that includes:

- 55 standard compute nodes with 64 hyperthreaded cores and 512GB of RAM
- 3 high-memory compute nodes with 48 cores and 1TB of RAM
- GPU nodes with Nvidia tesla k40, tesla k20, and titan GPUs
- A high-speed ethernet interconnect between 10 and 40 Gb/s

We used both the CPUs and all GPUs. All computation was written in Python 3 with associated packages. All code and models can be found at: https://github.com/UCDenver-ccp/Concept-Recognition-as-Translation.

### Span Detection

Span detection is the first task of our two-part approach to concept recognition. Even though there is quite a lot of previous work on this task, we aimed not only to explore the state-of-the-art methods (using language models), but also to find low-resource methods (using a CRF) and explore whether we can exploit hyperparameter tuning of simpler methods for more complex ones that build on the simple methods (using BiLSTMs). Thus, the goal is to explore the performance and resources of six canonical span detection algorithms [15, 44, 61] using the CRAFT shared task framework [51]. The underlying target representation for all algorithms are BIO tags [10], which label each word in the sequence as beginning (B), inside (I), or outside (O) an ontological concept mention; for example, the BIO tagging for the text mentions “red blood cells” and “white blood cells” in the phrase “red and white blood cells” can be seen in Table 3.

**Table 3.**
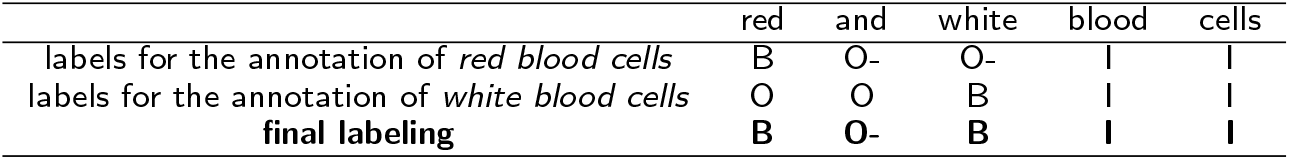
BIO labeling for the discontinuous and overlapping ontology class mentions in the phrase “red and white blood cells” (from PMCID:15314655). (The O-would simply be O in the canonical BIO labeling.)

|  | red | and | white | blood | cells |
| --- | --- | --- | --- | --- | --- |
| labels for the annotation of *red blood cells* | B | O- | O- | I | I |
| labels for the annotation of *white blood cells* | O | O | B | I | I |
| **final labeling** | **B** | **O-** | **B** | **I** | **I** |

Discontinuous annotations and overlapping annotations [17] make BIO labeling challenging to define formally. Discontinuous annotations are annotations composed of two or more non-continuous text spans. The difficulty in translation is how to label the intervening text between the two discontinuous spans, *i.e.*, as I (inside) or O (outside). Since discontinuous annotations are rare (no more than 7% of the total words in all concept mentions (see Table 4), many previous systems ignore them (*e.g.*, [4, 6]). Here we introduce and evaluate a novel approach simpler than previous work (*e.g.*, [16, 17]) to represent such annotations. Discontinuous annotations contain fewer words within their component text spans compared to the words between the spans, except for PR (see Table 4). Thus, we created a new label, O-, which signifies the text between the discontinuous spans of these annotations (see Table 3 for an example of “O-”). We refer to this expanded set of BIO tags as BIO-tags.

**Table 4.**
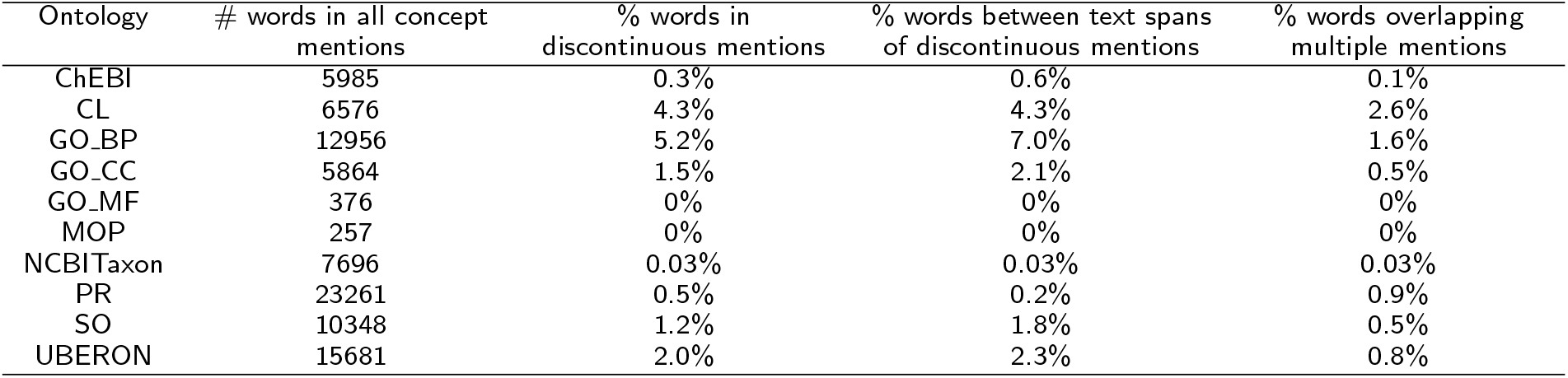
Quantification of discontinuous and overlapping words in all concept mentions. All numbers are based on the number of words, not concepts.

Overlapping spans create a different problem: the possibility of multiple labels for a word in the sequence, only one of which can persist for training purposes. For example “red”, “and”, and “white” all have two conflicting labels depending on the concept mention labeled (see Table 3). To address this problem, we prioritize the beginnings of concept mentions (B) to capture as many concepts as possible, even if some words in the multi-word concepts will not formally be captured. In our example then, the annotations based on the BIO-tags would be “red…” and “white blood cell” separately. Other approaches (*e.g.*, [62]) rewrite texts with conjunctions to unwind them, but that would not allow us to use the shared task framework, which depends on the unaltered text for evaluation.

Before training models, CRAFT is preprocessed into a word-tokenized BIO-tag format using the methods described above. This preprocessed data is used to train, tune, and evaluate all span detection algorithms. The input to each algorithm is a sentence as a sequence of words represented as word features, word embeddings, character embeddings, or language-model-contextualized word embeddings that are then mapped to a sequence of BIO-tags as the output. Each algorithm and its corresponding input representation is described in more detail below, focusing on the resources needed, the parameters to tune, and the final training framework used.

#### Conditional Random Fields (CRFs)

CRFs are of interest because they use the least computational resources of any of the approaches we have implemented and can generally be easily trained on contemporary laptops. A CRF is a discriminative algorithm that utilizes a combination of arbitrary, overlapping and agglomerative observation features from both the past and future to predict the output sequence [11]. The words in a sentence are the input sequence along with features for each word including the case of the word, the three words before the word for added context, its part-of-speech tag, and the part-of-speech tags for the two words ahead of the word of interest. The output sequence is a BIO-tag for each word. The output sequences are directly connected to the inputs via the sentence- and word-level features. To optimize F1 score for each ontology, we tuned the CRF by conducting a randomized search of the hyperparameter space of both the L1 and L2 regularization penalties, with 3-fold cross-validation, which guarantees that the global optimum is found [11]. With these optimal parameters, we evaluated the model using 5-fold cross-validation.

#### Bidirectional Long Short Term Memory (BiLSTM)

BiLSTMs are of interest because much prior work uses and builds on them with significant effort and resources put into tuning. Instead of tuning more complex systems, we aimed to test whether the parameters for the simplest model could generalize to more complex models built on the simple one. An LSTM is a special form of a recurrent neural network that by default remembers information for long periods of time, allowing for more distant context to be used by the algorithm [12]. It is a chain of memory cells stitched together to allow long-term and short-term memory. Each memory cell contains four neural networks, including a forget gate (information to drop), a new information gate (information to add), and an output gate to the next cell (information to propagate). The LSTM architecture lends itself to sequence-to-sequence tasks such as ours, in which the input is a sequence of words and the output is the corresponding sequence of BIO-tags. However, the inputs must be vectors, so we use an embedding layer that maps each word in the training data to a fixed-length vector to create the semantic vector space of the training data. Due to varying lengths of sentences, we padded all input sequences and output sequences to the maximum number of words among all sentences (approximately 400). Additionally, since the context for a word can be before or after the word itself, we used Bidirectional LSTMs (BiLSTMs), which first run through the network forward through the sequence and then backwards, thereby allowing the usage of the context on either side of a word [13].

Tuning a BiLSTM is resource-intensive, so we aimed to test whether the parameters tuned on a simple BiLSTM could translate across more complex BiLSTM models. Therefore, to conserve resources, hyperparameters tuned here are used for all other more complex BiLSTM approaches: BiLSTM-CRF, char-embeddings, and BiLSTM-ELMo. The hyperparameter search attempts to optimize F1 score; however, unlike the CRF setting, there are no guarantees of finding optimal parameters. The large hyperparameter space and the high cost of each evaluation are barriers to identifying optimal values. We followed established heuristics for tuning a BiL-STM [63], using GPUs to speed up evaluation. The four main hyperparameters to tune are the optimizer, the batch size, the number of epochs, and the number of neurons or hidden units [63]. The classic optimizer is the stochastic gradient descent (SGD), but newer approaches have proven less variant and faster [64], including RMSProp [65] and Adam [66]. For named entity recognition tasks, many have used SGD [30–32, 34–36, 38, 40], and a smaller number have used RMSProp [39, 41] and Adam [37]. We chose to focus on RMSProp (Root Mean Square Propagation), which updates the learning rate for each parameter separately and automatically using the exponential average of the square of the gradient in order to weight the more recent gradient updates over the less recent ones [65]. Thus, the learning rate requires very little tuning, and so we chose not to change the default learning rate of 0.0001. Additionally, it implicitly performs simulated annealing in that it automatically decreases the size of the gradient step if it is too large, so as not to overshoot the minima.

Next, multiple time-consuming experiments of different combinations of batch sizes, epochs, and neurons are needed to find the best parameters since they are interrelated. The batch size determines the number of examples to run together to help speed up the time-consuming training process. The larger the batch, the faster the runs, but the less nuanced the results. Typical batches range from 1 to 64 and thus we tested 18, 36, 53, and 106 (all of which cleanly divide a validation set of 10% of the training data). The epochs are the number of repeat experiments to run, as the LSTM can result in very different results based on each random initial condition. The larger the number of epochs, the more time the LSTM takes to run. Due to our limited memory and time, we tested a small (10) and larger (100) number of epochs. For all runs, 10% of the data was used for validation for each epoch, as well as overall. Lastly, to determine the number of neurons or the hidden states, we utilized this formula [67]:

*N*_*h*_ = 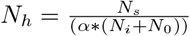
*N*_*i*_ = number of input neurons.
*N*_0_ = number of output neurons.
*N*_*s*_ = number of samples in training data set.
*α* = an arbitrary scaling factor, usually between 2 and 10.

We found that varying *α* between 2 and 10 yields between 3 and 12 neurons, and thus we took the two extremes of 3 and 12 to test.

Overall, we conducted 16 experiments (one optimizer, four batch sizes, two epoch sizes, and two neuron sizes) per ontology to find the optimal hyperparameters for the BiLSTM for each ontology. F1 score cannot be optimized directly with an LSTM; instead, the models aim to minimize errors. We chose the categorical cross-entropy as the loss function, as it is the default loss function for multi-class classification problems such as ours, in which four categories are used for BIO-tagging.

#### BiLSTM combined with CRF (BiLSTM-CRF)

A BiLSTM-CRF is the architecture of a regular BiLSTM with a CRF as the last layer [30]. The BiLSTM provides the feature weights for the CRF, which provides sequence-level features. The tuning processes would be exactly the same as for the previously discussed BiLSTM; however, to determine if the simple BiLSTM parameters can be used in a more complex model, we used the same tuned hyperparameters for each ontology found for the BiLSTM and then added the CRF layer on top. This is a simple extension of the previous model, which can be trained on CPUs since the BiLSTM tuning is already done.

#### BiLSTM with character embeddings (Char-Embeddings)

The BiLSTM and BiLSTM-CRF approaches use word embeddings for all of the words in the training data; thus, any word not in the training set cannot be translated to a BIO-tag and will be unknown. To combat this, we tried a different underlying sequence representation based on the characters to create character embeddings for each word. As each word is a sequence of characters, this approach can create a representation for any unknown words using these character embeddings. Again, to determine if the simple BiLSTM parameters can be used in more complex models, we utilized the same parameters as the tuned BiLSTM. In terms of resources, training the character embeddings adds a significant amount of time and memory using CPUs.

#### BiLSTM and Embeddings from a Language Model (BiLSTM-ELMo)

BiLSTM-ELMo is a BiLSTM with a new underlying sequence representation from the language model ELMo [44]. The original ELMo is a language model trained on the 1 Billion Word Benchmark set, which includes approximately 800 million tokens of news crawl data from the general-domain WMT 2011. ELMo representations are contextual (with modeling of word polysemy), deep (as the BiLSTM is pretrained on a large text corpus), and character-based (thus allowing for representations of words unseen in training). Again, to determine if the simple BiLSTM parameters can be used in more complex models, we wanted to use the already tuned parameters from our original BiLSTM, but due to limited resources, we ran out of memory quite quickly in training the BiLSTM-ELMo. Experimenting with the same aforementioned batch sizes, we found that a batch size of 18 could run for all ontologies. Thus, we took the optimal hyperparameters with a batch size of 18 for each ontology.

#### Bidirectional Encoder Representations from Transformers for Biomedical Text Mining (BioBERT)

BioBERT is a biomedical-specific language model pre-trained on biomedical documents from both PubMed abstracts (PubMed) and PubMed Central full-text articles (PMC) based on the original BERT architecture [15]. Briefly, BERT is a contextualized word representation model pre-trained using bidirectional transformers. It then uses a masked language model to predict randomly masked words in a sequence from the full context on either sides of the word (instead of scanning one direction at a time), creating bidirectional representations. BioBERT adds the additional layer of biomedical specific training data, as it is known that general-domain-trained algorithms do not usually perform well in the biomedical field [68, 69]. We chose to use the BioBERT + PubMed + PMC model because it is the most similar to CRAFT, the articles of which all appear in PMC. Due to the generalizabilty of BERT and BioBERT, they require minimal fine-tuning to utilize for other tasks, especially for this task since the documents in CRAFT are most likely included in the PMC training data. Thus, we utilized the default fine-tuning parameters for named entity recognition, including a learning rate of 1 *×* 10^−5^, a training batch size of 32, evaluation and prediction batch sizes of 8 each, and 10 training epochs [15], which requires minimal resources aside from a GPU. Similar to the LSTM models, validation is done within each epoch.

### Concept Normalization

The final step in the concept recognition task is concept normalization, *i.e.*, the normalization of the detected spans or text mentions of concepts to their respective unique ontology class identifiers (class IDs). For example, the text mention “white blood cell” is normalized to the class ID CL:0000738 and the text mention “red … blood cell” to the class ID CL:0000232 (see Figure 1). To the best of our knowledge, this is the first attempt to reframe and explore this task as a translation problem by translating the characters of all the text mentions to the characters of the ontology class identifiers. Usually, for translation from one language to another, there is an assumed underlying structure and semantics of both languages that is captured at least in part by any algorithm that aims to automate the translation process. For this task, the input in the form of English text mentions contains the structure and semantics of the English language [70], as well as the rich history of its development [71, 72]. On the other hand, the unique numeric class IDs that these concept mentions are annotated to supposedly contain no structure or semantics [73]. Thus, from the outset, one would not expect translation from text mentions to class IDs to work, yet our results suggest that it does. We designed a series of experiments to understand what signals in the ontology class IDs are important to the performance. To implement the idea of concept normalization as machine translation, we use the popular Open Neural Machine Translation system (OpenNMT) [18]. It implements stacked BiLSTMs with attention models and learns condensed vector representations of characters from the training data, processing one character at a time. One layer of the sequence-to-sequence LSTM model includes four main components: an encoder, a decoder, an attention mechanism, and a softmax layer. There are stacks of multiple layers of encoders, attention mechanisms, and decoders before the softmax layer at the top. The input to OpenNMT is the sequence of characters for the text mentions (*e.g.*, “w h i t e b l o o d c e l l”, with each character separated by a space to show it is a sequence of characters and not words). The size of this input sequence for the encoder is the length of the longest text mention in the training data by character count, which could be from 1 to 1000 characters depending on the ontology (*e.g.*, “white blood cell” has 16 characters including the spaces between words). Any input that is shorter than the maximum character length is padded at the end with null characters. Then for each text mention, the output is the class ID, similarly in the form of characters (*e.g.*, for the input “w h i t e b l o o d c e l l”, the output is “CL : 0000738” including the ontology namespace and the colon). Analogously, the output size is the maximum number of characters among the ontology class IDs, with added null characters if the sequence is shorter (*e.g.*, “C L : 0 0 0 0 7 3 8” has 10 characters). The maximum number of characters in the class IDs range among the ontologies from 7 to 20 characters in the core annotation sets and 10 to 83 in the core+extensions sets. This discrepancy arises from the naming convention of extension classes with both the additional “_EXT” in the namespace along with the textual class IDs, as detailed in the materials for CRAFT. We used the default parameter settings to begin to explore this idea as this approach is resource-intensive, requiring large amounts of memory and CPUs.

To train OpenNMT, pairs of text mentions and class IDs are required. For more generalized training, in addition to the linked text mentions and class IDs of the CRAFT concept annotations, we also used as training data the primary labels and synonyms (extracted from the .obo files distributed as part of the corpus) of classes not used for annotation in the corpus simply due to the fact that they are not mentioned in the articles of the corpus. For each of these classes, we used the name of the class and its synonyms as the text mentions that map to it; for example, CL:0000019, the Cell Ontology class representing sperm cells, does not occur in the CRAFT training data, so we added to the training data its primary label “sperm” and its exact synonyms “sperm cell”, “spermatozoon”, and “spermatozoid” as quasi-mentions that map to this ontology class. (See Table 2 for counts of ontology classes whose primary labels and synonyms were added to the training data.) Note that CRAFT annotators curated lists of unused classes that either were too difficult to reliably annotate with and/or for which extension classes were alternatively created or used; the labels and synonyms of these classes were not added as training data. By adding these metadata of these classes from the .obo files, we not only add a significant amount of training data (amounting to thousands of more classes per ontology), but also ensure that all current ontology classes are captured by OpenNMT (with the exception of those purposefully not used by the CRAFT curators).

With all of these concepts, training sets can be assembled at the type level (for which there is one mapping of a given text mention to a class ID regardless of frequency) or the token level (for which all mappings of text mentions to class IDs are included, even though some are the same string and class ID, only occurring in different places in the text). Training and testing with tokens takes into account the frequency of occurrence in the corpus (token-ids); using types ignores frequency of occurrence in training and evaluation (type-ids). The token-ids are used for the full end-to-end system as it captures all the data including frequency. However we compare token-ids to type-ids as well to determine if there is a performance difference, and type-ids are used for some experiments to better understand how concept normalization as machine translation works. We also explored the performance with and without the extension classes. The extension classes greatly increase the size of the training data, and have somewhat different performance characteristics. Tuning was over a 90-10 data split for training to validation over all tokens, and default training parameter settings were used.

As framing concept normalization as a machine translation problem is unconventional, we aimed to explore how this approach might be exploiting semantic information in the ontology class IDs (the core set only) used as output by transforming them in various ways to see how performance changes. To start, the type-ids are compared to the token-ids, and going forward all further experiments use the type-ids, as they are a smaller set and thus faster to run. If the frequency of the text mention and class ID mattered, then we would see a drop in performance from token-ids to type-ids. The next approach was to use the same IDs but scramble the relationship between text mention and class ID (“shuffled-ids”). Another was to replace the class IDs with random numbers of the same length, drawn without replacement as to have no repeats (“random-ids”). If there were information in the specific class IDs, we would expect to see a drop in performance for shuffled-ids relative to type-ids. If there were information in the distribution of class IDs but not in specific ones, then we would expect a further drop in performance for random-ids relative to shuffled-ids. We further tested to see if we could add information to the class IDs by alphabetizing them by the text mention and assigning consecutive IDs (“alphabetical-ids”). Text mentions that have similar prefixes (*e.g.*, proteins BRCA1 and BRCA1 C-terminus-associated protein) would have consecutive alphabetical IDs (PR:089212 and PR:089213, respectively), potentially giving the sequence-to-sequence learner additional information. For all experiments, we maintained the ontology prefixes before the unique number identifiers and only changed the numbers (as seen in the example above for alphabetical-ids). However, not all text mentions map to class IDs solely with numbers. For example, the class ID for the text mention “phylum” is NCBITaxon:phylum. These types of text mentions and class IDs are rare in ontologies in the core set and thus were not changed in any experiments. In the evaluation of alternative output targets, we calculate both exact matches between ontology class identifiers overall and on a per-character basis, as OpenNMT translates per character. Furthermore, we conducted error analyses of these runs to better understand what underlies the translation and suggest future improvements.

## Results

Overall we achieve near- or above-state-of-the-art performance on the concept annotation task of the CRAFT Shared Tasks, with direct comparison to Furrer et al. [4] on the full end-to-end system using the corresponding evaluation platform as described above (see Tables 5 and 6). Recall that partial credit is awarded both for span detection and concept normalization, so we also evaluate each separately to understand what drives the performance in the full end-to-end system. Even with the state-of-the-art performance, there is still room for improvement for all ontologies in the core and core+extensions sets, especially for PR and PR_EXT, respectively.

**Table 5.** Full end-to-end system evaluation on the core set comparing F1 score. For all columns, the span detection algorithm is listed and it is assumed that OpenNMT is the concept normalization algorithm. UZH@CRAFT-ST is the best system for each ontology from Furrer et al. [4]. The best algorithm is bolded with an asterisks*.

| Ontology | CRF | BiLSTM | BiLSTM-CRF | Char-Embeddings | BiLSTM-ELMo | BioBERT | UZH@CRAFT-ST |
| --- | --- | --- | --- | --- | --- | --- | --- |
| ChEBI | 0.7882 | 0.6394 | 0.5027 | 0.5942 | 0.0550 | **0.7885\*** | 0.7700 |
| CL | 0.6779 | 0.5134 | 0.3859 | 0.5611 | 0.0526 | **0.6994\*** | 0.6657 |
| GO_BP | 0.7505 | 0.5137 | 0.3642 | 0.6182 | 0.0720 | 0.7405 | **0.8037\*** |
| GO_CC | 0.7225 | 0.1689 | 0.3049 | 0.3244 | 0.0506 | **0.7762\*** | 0.7645 |
| GO_MF | 0.9778 | 0.9770 | 0.9778 | 0.8906 | 0.3704 | 0.9783 | **0.9838\*** |
| MOP | 0.8129 | 0.7721 | 0.7158 | 0.5985 | 0.0930 | **0.8742\*** | 0.8705 |
| NCBITaxon | 0.9026 | 0.7736 | 0.8391 | 0.8518 | 0.0948 | 0.8910 | **0.9694\*** |
| PR | 0.4040 | 0.3136 | 0.2827 | 0.2732 | 0.0516 | 0.5295 | **0.8026\*** |
| SO | 0.8987 | 0.4106 | 0.4096 | 0.7815 | 0.0813 | **0.9054\*** | 0.9027 |
| UBERON | 0.7474 | 0.6812 | 0.5029 | 0.6901 | 0.0793 | **0.7670\*** | 0.7488 |

**Table 6.** Full end-to-end system evaluation on the core+extensions set comparing F1 score for the top two algorithms found in the core set. For all columns, the span detection algorithm is listed and it is assumed that OpenNMT is the concept normalization algorithm. UZH@CRAFT-ST is the best system for each ontology from Furrer et al. [4]. The best algorithm is bolded with an asterisk*.

| Ontology | CRF | BioBERT | UZH@CRAFT-ST |
| --- | --- | --- | --- |
| ChEBI_EXT | 0.7891 | 0.8039 | **0.8209\*** |
| CL_EXT | 0.7381 | **0.7491\*** | 0.7484 |
| GO_BP_EXT | 0.7279 | 0.7353 | **0.8138\*** |
| GO_CC_EXT | 0.8738 | **0.8983\*** | 0.8936 |
| GO_MF_EXT | 0.6413 | 0.6255 | **0.7438\*** |
| MOP_EXT | 0.8000 | **0.8651\*** | 0.8437 |
| NCBITaxon_EXT | 0.8710 | 0.8624 | **0.9722\*** |
| PR_EXT | 0.4397 | 0.5188 | **0.8011\*** |
| SO_EXT | 0.7682 | 0.7829 | **0.9187\*** |
| UBERON_EXT | 0.7558 | 0.7711 | **0.7714\*** |

Not surprisingly, BioBERT outperforms almost all other span detection algorithms, except for the CRF (for GO_BP, NCBITaxon, GO_MF_EXT, and NCBITaxon_EXT). (See Tables 5 and 6 for full results as compared to those of Furrer et al. [4] (UZH@CRAFT-ST).) The BiLSTMs overall did not perform the best, but it does seem that reusing the simplest model parameters in the more complex models of BiLSTM-CRF and a BiLSTM with character embeddings (Char-Embeddings) either maintains the same performance or sometimes increases performance, especially for the Char-Embeddings model. However, these simple model parameters cannot be reused for the BiLSTM-ELMo model. In comparison to UZH@CRAFT-ST, for the core set, ChEBI, CL, GO_CC, MOP, SO, and UBERON modestly outperform the best system, whereas CL_EXT, GO_CC_EXT, and MOP_EXT just barely outperformed the best system for the core+extensions set. Even for the ontologies that are lower than the best system, they are usually in close proximity (within 0.10). Note that GO_MF has the highest performance of all ontologies, most likely due to the very few annotation classes included in the ontology (see Tables 1 and 2). Lastly, for some ontology annotation sets (specifically, ChEBI, CL, GO_CC, UBERON, CL_EXT, GO_MF_EXT, and UBERON_EXT), all systems (including UZH@CRAFT-ST) perform less competently, with F1 scores below 0.80.

It does appear then that translation is a salient avenue to explore for a purely machine learning approach to concept recognition, which at least for the core set is comparable to the state of the art. We also describe the resources needed for each algorithm (see Table 7); and the tuning, training and evaluation of the span detection algorithms (see Tables 8–14), and the concept normalization algorithm (see Tables 15–22).

**Table 7.** Hardware, memory, and time used for training for all evaluated algorithms. A given training time specifies the total hours if training for all ontology annotation sets were run consecutively, but these can be parallelized by ontology. *Parallelized per ontology due to time constraints. **Runs significantly faster on GPUs. ***Total free RAM available. ConceptMapper runs on CPUs but has no training, as it is a dictionary-based lookup tool, hence the specifications as N/A.

| Algorithm | Hardware | Training Memory (GBs) | Training Time (hrs) |
| --- | --- | --- | --- |
| CRF | CPUs | 2-13 | 1-4 |
| BiLSTM* | GPUs/CPUs** | 17 | 29 |
| BiLSTM-CRF | CPUs | 7 | 15 |
| Char-Embeddings | CPUs | 30 | 84 |
| BiLSTM-ELMo* | GPUs | 42 | 700-1000 |
| BioBERT | GPUs/CPUs** | 5 | 20 |
| UZH@CRAFT-ST BioBERT* [4] | GPUS | 120*** | 200 |
| OpenNMT* | CPUs | 620 | 515 |
| ConceptMapper [20] | CPUs | N/A | N/A |

**Table 8.** Span detection F1 score results for all algorithms tested against the core evaluation annotation set for the 30 held-out articles. The best-performing algorithm per ontology is bolded with an asterisk*.

| Ontology | CRF | BiLSTM | BiLSTM-CRF | Char-Embeddings | BiLSTM-ELMo | BioBERT |
| --- | --- | --- | --- | --- | --- | --- |
| ChEBI | 0.7234 | 0.6545 | 0.5000 | 0.5280 | 0.0620 | **0.9091\*** |
| CL | 0.8333 | 0.5882 | 0.3774 | 0.8000 | 0.0000 | **0.9231\*** |
| GO_BP | **0.8677\*** | 0.5498 | 0.3661 | 0.6346 | 0.0685 | 0.8646 |
| GO_CC | 0.9412 | 0.1379 | 0.2689 | 0.2581 | 0.1000 | **0.9444\*** |
| GO_MF | >**0.9999\*** | >**0.9999\*** | >**0.9999\*** | 0.8421 | 0.0000 | >**0.9999\*** |
| MOP | >**0.9999\*** | >**0.9999\*** | >**0.9999\*** | >**0.9999\*** | 0.0000 | >**0.9999\*** |
| NCBITaxon | **0.9959\*** | 0.8551 | 0.9440 | 0.9569 | 0.0711 | 0.9453 |
| PR | 0.4351 | 0.2979 | 0.2151 | 0.0995 | 0.0339 | **0.8199\*** |
| SO | **0.9435\*** | 0.4935 | 0.4897 | 0.8203 | 0.1059 | 0.9081 |
| UBERON | 0.7913 | 0.7206 | 0.4758 | 0.7440 | 0.0854 | **0.8826\*** |

**Table 9.**
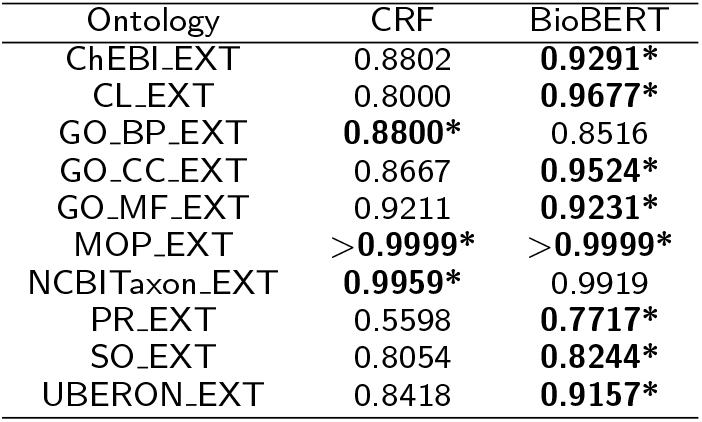
Span detection F1 score results for all algorithms tested against the core+extensions evaluation annotation set for the 30 held-out articles. The best-performing algorithm per ontology is bolded with an asterisk*.

| Ontology | CRF | BioBERT |
| --- | --- | --- |
| ChEBI_EXT | 0.8802 | **0.9291\*** |
| CL_EXT | 0.8000 | **0.9677\*** |
| GO_BP_EXT | **0.8800\*** | 0.8516 |
| GO_CC_EXT | 0.8667 | **0.9524\*** |
| GO_MF_EXT | 0.9211 | **0.9231\*** |
| MOP_EXT | >**0.9999\*** | >**0.9999\*** |
| NCBITaxon_EXT | **0.9959\*** | 0.9919 |
| PR_EXT | 0.5598 | **0.7717\*** |
| SO_EXT | 0.8054 | **0.8244\*** |
| UBERON_EXT | 0.8418 | **0.9157\*** |

**Table 10.** F1 score results for detection of discontinuous spans for all algorithms tested against the core evaluation annotation set for the 30 held-out articles. (Note that there are no discontinuous spans the in the GO_MF, MOP, and NCBITaxon sets.)

| Ontology | Support | CRF | BiLSTM | BiLSTM-CRF | Char-Embeddings | BiLSTM-ELMo | BioBERT |
| --- | --- | --- | --- | --- | --- | --- | --- |
| ChEBI | 14 | 0 | 0 | 0 | 0 | 0 | 0 |
| CL | 175 | 0.1176 | 0.1158 | 0.1171 | 0 | 0.0107 | 0.1818 |
| GO_BP | 272 | 0.0952 | 0 | 0.014 | 0 | 0.007 | 0.2742 |
| GO_CC | 14 | 0.1053 | 0 | 0 | 0.0526 | 0 | 0.3 |
| PR | 44 | 0 | 0 | 0 | 0 | 0 | 0.08 |
| SO | 45 | 0.0408 | 0 | 0 | 0 | 0 | 0.3 |
| UBERON | 118 | 0.04 | 0 | 0 | 0 | 0 | 0.0915 |

**Table 11.** F1 score results for detection of discontinuous spans for all algorithms tested against the core+extensions evaluation annotation set for the 30 held-out documents. (Note that there are no discontinuous spans in the MOP_EXT and NCBITaxon_EXT sets.)

| Ontology | Support | CRF | BioBERT |
| --- | --- | --- | --- |
| ChEBI_EXT | 19 | 0 | 0 |
| CL_EXT | 175 | 0.1164 | 0.1608 |
| GO_BP_EXT | 287 | 0.085 | 0.2651 |
| GO_CC_EXT | 30 | 0.1579 | 0.3721 |
| GO_MF_EXT | 20 | 0 | 0 |
| PR_EXT | 44 | 0 | 0.1224 |
| SO_EXT | 72 | 0.1505 | 0.2979 |
| UBERON_EXT | 133 | 0.0485 | 0.2128 |

**Table 12.** CRF tuning parameters and results. The overall memory usage for all tuning was 6 GB.

| Ontology | L1 | L2 | Time (hours) | F1 Score (macro) | F1 Score (micro) |
| --- | --- | --- | --- | --- | --- |
| ChEBI | 0.0862 | 0.000186 | 3 | 0.61 | 0.99 |
| CL | 0.00477 | 0.0473 | 3 | 0.73 | >0.99 |
| GO_BP | 0.0862 | 0.000186 | 3.17 | 0.63 | 0.99 |
| GO_CC | 0.269 | 0.00892 | 3 | 0.55 | 0.99 |
| GO_MF | 0.215 | 0.00392 | 3 | 0.66 | 0.99 |
| MOP | 0.0862 | 0.000186 | 3 | 0.65 | >0.99 |
| NCBITaxon | 0.0862 | 0.000186 | 3 | 0.61 | >0.99 |
| PR | 0.00477 | 0.0473 | 3.25 | 0.54 | 0.97 |
| SO | 0.315 | 0.00578 | 3.22 | 0.66 | >0.99 |
| UBERON | 0.221 | 0.003005 | 3.3 | 0.63 | 0.99 |

**Table 13.** BiLSTM tuning parameters and results that are used for BiLSTM-CRF and Char-Embeddings also.

| Ontology | Batch Size | # Epochs | # Neurons | Time (hours) | Memory (GBs) | F1 Score (macro) | F1 Score (micro) |
| --- | --- | --- | --- | --- | --- | --- | --- |
| ChEBI | 53 | 10 | 12 | 99 | 6.5 | 0.67 | >0.99 |
| CL | 36 | 10 | 12 | 92 | 6.5 | 0.74 | >0.99 |
| GO_BP | 36 | 10 | 12 | 99 | 6.5 | 0.68 | >0.99 |
| GO_CC | 106 | 100 | 12 | 97 | 6.5 | 0.66 | >0.99 |
| GO_MF | 106 | 10 | 3 | 108 | 8.4 | 0.99 | >0.99 |
| MOP | 106 | 10 | 12 | 99 | 6.4 | 0.61 | >0.99 |
| NCBITaxon | 106 | 100 | 12 | 95 | 6.5 | 0.96 | >0.99 |
| PR | 36 | 10 | 12 | 95 | 6.5 | 0.71 | >0.99 |
| SO | 36 | 100 | 12 | 98 | 6.5 | 0.66 | >0.99 |
| UBERON | 18 | 10 | 12 | 97 | 6.5 | 0.68 | >0.99 |

**Table 14.** BiLSTM-ELMo parameters and results. Due to limited resources, the batch size is 18 for all ontologies.

| Ontology | Batch size | # Epochs | # Neurons | F1 Score (macro) | F1 Score (micro) |
| --- | --- | --- | --- | --- | --- |
| ChEBI | 18 | 10 | 3 | 0.65 | >0.99 |
| CL | 18 | 10 | 12 | 0.72 | >0.99 |
| GO_BP | 18 | 10 | 12 | 0.66 | >0.99 |
| GO_CC | 18 | 10 | 3 | 0.65 | >0.99 |
| GO_MF | 18 | 100 | 3 | 0.66 | >0.99 |
| MOP | 18 | 100 | 12 | 0.61 | >0.99 |
| NCBITaxon | 18 | 100 | 12 | 0.96 | >0.99 |
| PR | 18 | 10 | 12 | 0.71 | >0.99 |
| SO | 18 | 100 | 12 | 0.65 | >0.99 |
| UBERON | 18 | 10 | 12 | 0.68 | >0.99 |

**Table 15.**
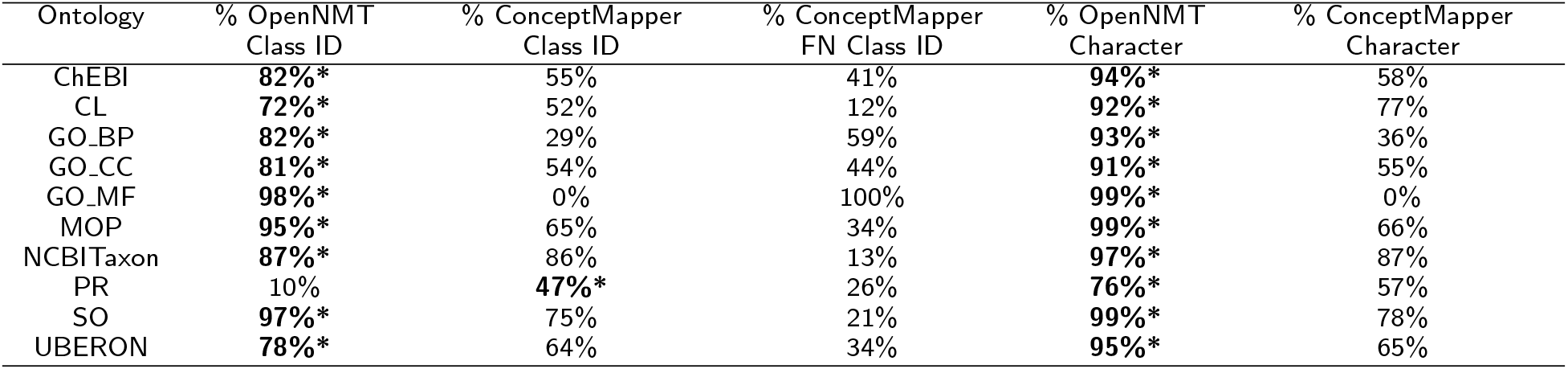
Concept normalization exact match results on the core evaluation annotation set for the 30 held-out documents compared to the baseline ConceptMapper approach. We report both the percent exact match on the class ID level and the character level. We also report the percentage of false negatives (FN) for ConceptMapper (*i.e.* no class ID prediction for a given text mention). Note that the best performance between OpenNMT and ConceptMapper is bolded with an asterisk* for both class ID and character level.

| Ontology | % OpenNMT<br>Class ID | % ConceptMapper<br>Class ID | % ConceptMapper<br>FN Class ID | % OpenNMT<br>Character | % ConceptMapper<br>Character |
| --- | --- | --- | --- | --- | --- |
| ChEBI | **82%\*** | 55% | 41% | **94%\*** | 58% |
| CL | **72%\*** | 52% | 12% | **92%\*** | 77% |
| GO_BP | **82%\*** | 29% | 59% | **93%\*** | 36% |
| GO_CC | **81%\*** | 54% | 44% | **91%\*** | 55% |
| GO_MF | **98%\*** | 0% | 100% | **99%\*** | 0% |
| MOP | **95%\*** | 65% | 34% | **99%\*** | 66% |
| NCBITaxon | **87%\*** | 86% | 13% | **97%\*** | 87% |
| PR | 10% | **47%\*** | 26% | **76%\*** | 57% |
| SO | **97%\*** | 75% | 21% | **99%\*** | 78% |
| UBERON | **78%\*** | 64% | 34% | **95%\*** | 65% |

**Table 16.**
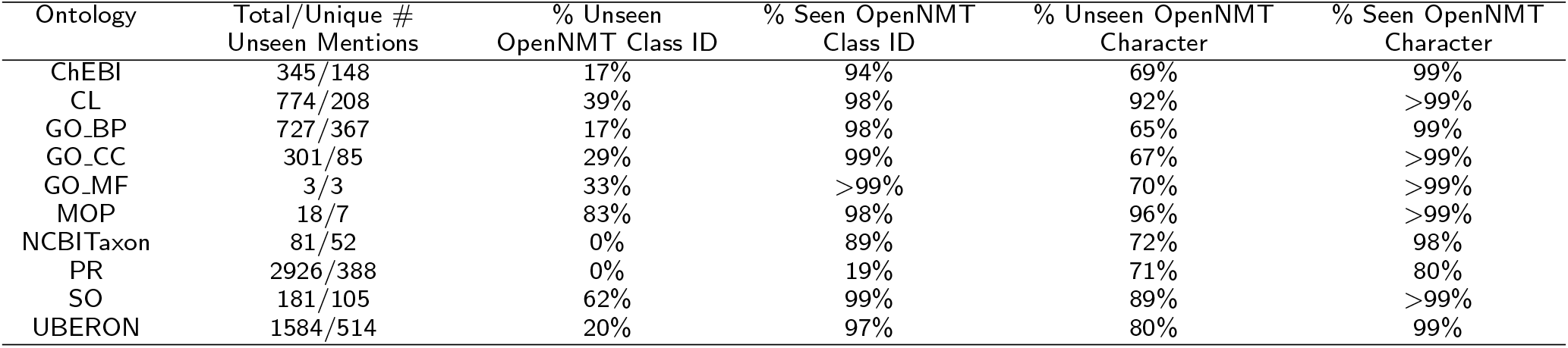
Exact match results for the unseen and seen text mentions (relative to the training data) for the core evaluation annotation set of the 30 held-out documents. Reporting the total number of unseen mentions/the number of unique unseen mentions along with the percent exact match on the class ID level and character level for both unseen and seen text mentions.

| Ontology | Total/Unique #<br>Unseen Mentions | % Unseen<br>OpenNMT Class ID | % Seen OpenNMT<br>Class ID | % Unseen OpenNMT<br>Character | % Seen OpenNMT<br>Character |
| --- | --- | --- | --- | --- | --- |
| ChEBI | 345/148 | 17% | 94% | 69% | 99% |
| CL | 774/208 | 39% | 98% | 92% | >99% |
| GO_BP | 727/367 | 17% | 98% | 65% | 99% |
| GO_CC | 301/85 | 29% | 99% | 67% | >99% |
| GO_MF | 3/3 | 33% | >99% | 70% | >99% |
| MOP | 18/7 | 83% | 98% | 96% | >99% |
| NCBITaxon | 81/52 | 0% | 89% | 72% | 98% |
| PR | 2926/388 | 0% | 19% | 71% | 80% |
| SO | 181/105 | 62% | 99% | 89% | >99% |
| UBERON | 1584/514 | 20% | 97% | 80% | 99% |

**Table 17.** Concept normalization exact match results on the core+extensions evaluation annotation set for the 30 held-out documents compared to the baseline ConceptMapper approach. We report both the percent exact match on the class ID level and the character level. We also report the percentage of false negatives (FN) for ConceptMapper (*i.e.* no class ID prediction for a given text mention). Note that the best performance between OpenNMT and ConceptMapper is bolded with an asterisk* for both class ID and character level.

| Ontology | % OpenNMT Class ID | % ConceptMapper Class ID | % ConceptMapper FN Class ID | % OpenNMT Character | % ConceptMapper Character |
| --- | --- | --- | --- | --- | --- |
| ChEBI_EXT | **86%\*** | 64% | 26% | **84%\*** | 66% |
| CL_EXT | **82%\*** | 67% | 11% | **93%\*** | 84% |
| GO_BP_EXT | **80%\*** | 34% | 44% | **76%\*** | 38% |
| GO_CC_EXT | **93%\*** | 80% | 18% | **94%\*** | 84% |
| GO_MF_EXT | **69%\*** | 60% | 30% | **69%\*** | 64% |
| MOP_EXT | **92%\*** | 64% | 35% | **97%\*** | 44% |
| NCBITaxon_EXT | 83% | **86%\*** | 13% | **93%\*** | 87% |
| PR_EXT | **15%\*** | 9% | 28% | **72%\*** | 21% |
| SO_EXT | **92%\*** | 19% | 40% | **91%\*** | 22% |
| UBERON_EXT | **81%\*** | 68% | 29% | **92%\*** | 75% |

**Table 18.** Exact match results for the unseen and seen text mentions (relative to the training data) for the core+extensions evaluation annotation set of the 30 held-out documents. Reporting the total number of unseen mentions/the number of unique unseen mentions along with the percent exact match on the class ID level and character level for both unseen and seen text mentions.

| Ontology | Total/Unique # Unseen Mentions | % Unseen OpenNMT Class ID | % Seen OpenNMT Class ID | % Unseen OpenNMT Character | % Seen OpenNMT Character |
| --- | --- | --- | --- | --- | --- |
| ChEBI_EXT | 476/188 | 32% | 92% | 67% | 85% |
| CL_EXT | 775/209 | 36% | 99% | 77% | 97% |
| GO_BP_EXT | 861/431 | 26% | 89% | 57% | 78% |
| GO_CC_EXT | 339/113 | 39% | 99% | 57% | 98% |
| GO_MF_EXT | 515/146 | 31% | 83% | 45% | 78% |
| MOP_EXT | 21/10 | 67% | 98% | 85% | >99% |
| NCBITaxon_EXT | 123/79 | 1% | 86% | 68% | 94% |
| PR_EXT | 3114/429 | 0% | 25% | 66% | 75% |
| SO_EXT | 318/183 | 51% | 94% | 69% | 92% |
| UBERON_EXT | 1609/532 | 23% | 96% | 79% | 95% |

**Table 19.** Percentage of non-existent class IDs out of the total number of mismatch class IDs for the core set for the training, validation and evaluation sets.

| Ontology | % Non-existent Class IDs in Training | % Non-existent Class IDs in Validation | % Non-existent Class IDs in Evaluation |
| --- | --- | --- | --- |
| ChEBI | 3% | 4% | 11% |
| CL | 0% | 0% | 0% |
| GO_BP | 2% | 2% | 2% |
| GO_CC | 2% | 4% | 1% |
| GO_MF | 1% | 1% | 50% |
| MOP | 0% | 2% | 0% |
| NCBITaxon | 7% | 7% | 11% |
| PR | 10% | 10% | 2% |
| SO | 0% | 1% | 0% |
| UBERON | 0% | 1% | 2% |

**Table 20.** Percentage of non-existent class IDs out of the total number of mismatch class IDs for the core+extensions set for the training, validation and evaluation sets.

| Ontology | % Non-existent Class IDs in Training | % Non-existent Class IDs in Validation | % Non-existent Class IDs in Evaluation |
| --- | --- | --- | --- |
| ChEBI_EXT | 8% | 8% | 17% |
| CL_EXT | 1% | 1% | 9% |
| GO_BP_EXT | 2% | 2% | 3% |
| GO_CC_EXT | 4% | 4% | 21% |
| GO_MF_EXT | 2% | 2% | 4% |
| MOP_EXT | 0% | 6% | 0% |
| NCBITaxon_EXT | 9% | 9% | 25% |
| PR_EXT | 7% | 7% | 5% |
| SO_EXT | 1% | 1% | 2% |
| UBERON_EXT | 1% | 1% | 1% |

**Table 21.**
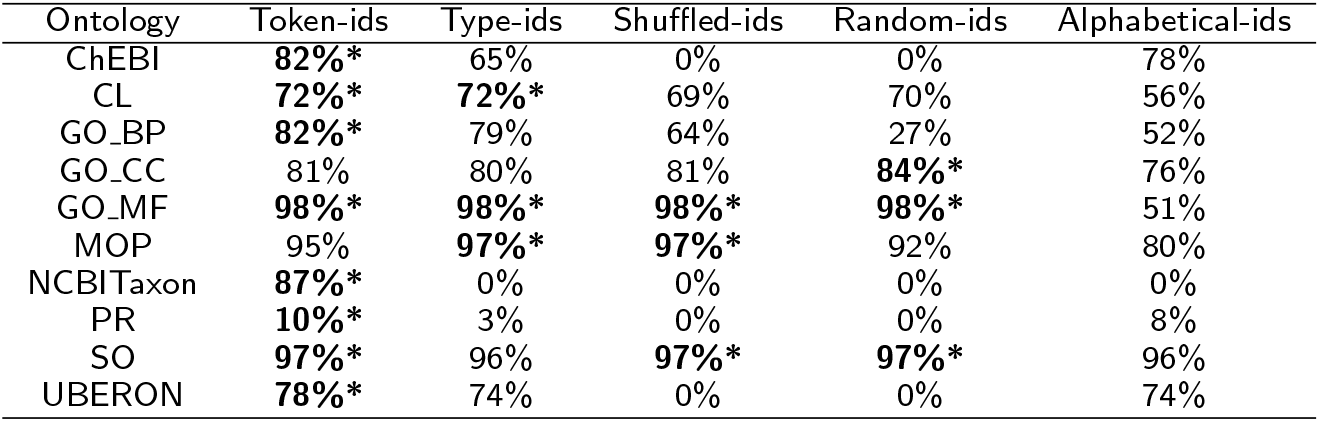
Exact match results for the concept normalization experiments on the core evaluation annotation set of 30 held-out documents. We report the exact match percentage on the class ID level. The highest percentage is bolded and with an asterisk*).

| Ontology | Token-ids | Type-ids | Shuffled-ids | Random-ids | Alphabetical-ids |
| --- | --- | --- | --- | --- | --- |
| ChEBI | **82%\*** | 65% | 0% | 0% | 78% |
| CL | **72%\*** | **72%\*** | 69% | 70% | 56% |
| GO_BP | **82%\*** | 79% | 64% | 27% | 52% |
| GO_CC | 81% | 80% | 81% | **84%\*** | 76% |
| GO_MF | **98%\*** | **98%\*** | **98%\*** | **98%\*** | 51% |
| MOP | 95% | **97%\*** | **97%\*** | 92% | 80% |
| NCBITaxon | **87%\*** | 0% | 0% | 0% | 0% |
| PR | **10%\*** | 3% | 0% | 0% | 8% |
| SO | **97%\*** | 96% | **97%\*** | **97%\*** | 96% |
| UBERON | **78%\*** | 74% | 0% | 0% | 74% |

**Table 22.** Exact match results for the concept normalization experiments on the core evaluation annotation set of 30 held-out documents. We report the exact match percentage on the character level. The highest percentage is bolded and with an asterisk*).

| Ontology | Token-ids | Type-ids | Shuffled-ids | Random-ids | Alphabetical-ids |
| --- | --- | --- | --- | --- | --- |
| ChEBI | **94%\*** | 89% | 60% | 58% | **94%\*** |
| CL | **92%\*** | **92%\*** | 86% | 80% | 65% |
| GO_BP | **93%\*** | 91% | 85% | 56% | 84% |
| GO_CC | 91% | **92%\*** | **92%\*** | 89% | 82% |
| GO_MF | **99%\*** | **99%\*** | **99%\*** | **99%\*** | 94% |
| MOP | 99% | **>99%\*** | 99% | 99% | 86% |
| NCBITaxon | **97%\*** | 74% | 73% | 68% | 74% |
| PR | **76%\*** | 75% | 40% | 30% | 74% |
| SO | **99%\*** | **99%\*** | **99%\*** | 98% | 96% |
| UBERON | **95%\*** | 93% | 69% | 54% | 88% |

**Table 23.** Abbreviations.

| Abbreviation | Meaning |
| --- | --- |
| NLP | Natural Language Processing |
| BioCreAtIve | Critical Assessment of Information Extraction in Biology |
| BioNLP-OST | BioNLP Open Shared Tasks |
| CRAFT | Colorado Richly Annotated Full-Text |
| CRAFT-ST | CRAFT Shared Task |
| OBO | Open Biomedical Ontologies |
| ChEBI | Chemical Entities of Biological Interest |
| CL | Cell Ontology |
| GO_BP | Gene Ontology Biological Processes |
| GO_CC | Gene Ontology Cellular Components |
| GO_MF | Gene Ontology Molecular Function |
| MOP | Molecular Process Ontology |
| NCBITaxon | NCBI Taxonomy |
| PR | Protein Ontology |
| SO | Sequence Ontology |
| UBERON | Uberon |
| EXT | Extension |
| BIO/BIO- tags | B: Beginning, I: Inside, O: Outside, O-: between a discontinuous span |
| CRF | Conditional Random Fields |
| BiLSTM/LSTM | Bidirectional Long Short Term Memory |
| RMSProp | Root Mean Square Propagation |
| ELMo | Embeddings from a Language Model |
| BERT | Bidirectional Encoder Representations from Transformers |
| BioBERT | Bidirectional Encoder Representations from Transformers for Biomedical Text Mining |
| OpenNMT | Open-source toolkit for Neural Machine Translation |
| CPU | Central Processing Unit |
| GPU | Graphics processing unit |
| OGER | OntoGene's Entity Recognizer |
| UZH@CRAFT-ST | University of Zurich at CRAFT Shared Task |
| CNN | Convolutional Neural Network |
| RNN | Recurrent Neural Network |
| Class ID/ID | Ontology class identifier |
| Token-ids | All mappings of spans to concepts; all unique strings and concepts in the text |
| Type-ids | One mapping of a given span to a concept regardless of frequency |
| Shuffled-ids | Shuffled the ontology class IDs |
| Random-ids | Assigned random ontology class IDs |
| Alphabetical-ids | Assigned new IDs based on alphabetizing the concept labels |
| core set | Annotations made with proper classes of the original given OBO |
| core+extensions set | Annotations to the core set and classes created as extensions of the ontologies |

### Training Resources

Training these algorithms requires significant computational resources due to the amount of data, tuning, and optimizing necessary. It is important to consider access to hardware, memory, and time when deciding on which algorithms to use for a task. Thus, here we report on those factors to aid other users in making these decisions (see Table 7). We report on these resources for the core annotation set only, as those for the core+extensions annotation set are similar.

Usually, having access to GPUs is helpful in speeding up training time as are hyperthreaded CPUs, since advanced hardware is quite important for some algorithms. Here, we began with CPUs and switched to GPUs as the time to train increased, as the latter will often speed up the processes significantly [74]. The CRF, BiLSTM-CRF, and Char-Embeddings ran in reasonable amounts of time on CPUs for all ontology annotation sets. For the BiLSTM itself though, we tuned it on GPUs and left it on GPUs once we had the optimized parameters. BioBERT also performed significantly faster on GPUs than on CPUs. At the same time, some algorithms need a GPU for practical running times due to the amount of tasks performed in it [74], such as the BiLSTM-ELMo model, where ELMo is the rate-limiting step.

All algorithms required a small amount of memory, except for OpenNMT, which required more than 600GB to train all ontologies. Due to its memory requirement, we parallelized it in terms of the ten ontologies, averaging around 60-80GBs per ontology. We also parallelized BiLSTMs and BiLSTM-ELMo due to similar considerations.

Choosing the right hardware with the right memory also hinges on the training time of the algorithm, which is particularly relevant for, *e.g.*, a shared task, where time is very important for meeting deadlines. We found that the fastest algorithm was the CRF, followed by BiLSTM-CRF and BioBERT. The UZH@CRAFT-ST also used BioBERT, and it took significantly longer than our BioBERT due to a differing number of epochs (55 and 10, respectively). The most time-consuming algorithms were BiLSTM-ELMo, with ELMo taking the majority of the time, and OpenNMT. In both cases though, similar to memory issues, we parallelized over the ten different ontologies to speed up the process. Still though, each ontology required 50-100 hours of supercomputer time.

### Span Detection Results

Overall, BioBERT and CRF perform best for span detection on both the core and core+extensions annotation sets (see Tables 8 and 9, respectively) as evaluated on the set of 30 held-out documents. Overall performance is very good, with all ontology F1 scores for both sets above 0.77 and most above 0.90. Note that the best results are seen for GO_MF, MOP, and MOP_EXT, most likely due to the fact that there are relatively few annotations in these sets (see Tables 1 and 2). One can see that BiLSTM-ELMo performs the worst and is most likely the reason the full end-to-end system is so poor for this algorithm. Even with some poor results, all algorithms can detect some discontinuous spans, and some discontinuous spans can be detected by at least one algorithm for all ontologies except for ChEBI, ChEBI_EXT, and GO_MF_EXT, which have among the fewest discontinuous spans (see Tables 10 and 11 for the core and core+extensions annotation sets, respectively). Due to the rarity of discontinuous spans, these tables suggest that we can just barely detect them and that more work is needed on these complex mentions specifically.

The most time-consuming and difficult part of this work is tuning the algorithms, whereas running and evaluating the models are generally quite fast. So that others can benefit from this work, we present the tuning results for the CRF and BiLSTM models, as well as the final parameters used for the BiLSTM-ELMo model. All tuning was parallelized in terms of the ten different ontologies for time and memory efficiency. We report both the macro- and micro-F1 scores because the “O” (outside) category for the BIO-tags appears significantly more often than any other tag. The micro-F1 scores take this into account and thus are significantly higher, whereas the macro-F1 score only looks at the raw scores, weighting each tag equally. Conversely, the macro-F1 scores are low due to the infrequent “O-” tag.

The CRF required the least amount of time for tuning (around three hours for each ontology) to find the optimal L1 and L2 regularization penalties (see Table 12). Of note is the fact that ChEBI, GO_BP, MOP, and NCBITaxon all share the same parameters and have very similar F1 scores. This may mean that these ontologies need more tuning with larger parameter spaces or that this signifies a universal parameter for most ontologies. In general, all macro-F1 scores are above 0.6 except for GO_CC and PR, which are still above 0.5. PR has the lowest score and CL has the highest, with room for improvement in both.

In contrast to the CRF tuning, tuning the BiLSTM required a significantly longer amount of time to achieve optimal performance on the training data. However, less time was required for the more complex BiLSTM algorithms (BiLSTM-CRF and Char-Embeddings) as we reused these parameters, even though it did not perform the best in the full-system evaluation. On average, training took around 98 hours per ontology, which is due in part to the number of parameters tested. That being said, each tested parameter value for batch size and numbers of epochs and neurons was required by at least one ontology for optimal performance. For batch size, the two most common were 106 (the largest value tested) and 36 (the second lowest value tested), found to be optimal for four ontologies each, while optimal batch sizes of 53 and 18 were found for ChEBI and UBERON, respectively. As for epochs, an optimal number of 10 was found for most of the ontologies, while an optimal number of 100 was found for GO_CC, NCBITaxon, and SO. For neurons, an optimal number of 12 was found for all ontologies except for GO_MF, for which an optimal number of 3 was found. About the same amount of memory (6.5 GB) was used for tuning all ontologies. In terms of optimization metrics, we calculated macro- and micro-F1 scores, as for the CRF model. In this case, all micro-F1 scores are above 0.99, while macro-F1 scores are equivalent to or greater than those for the CRF model, with all scores greater than 0.6, in contrast to the results for the final end-to-end system, for which the CRF model outperforms all BiLSTM algorithms. The highest F1 score is that for GO_MF and the lowest for MOP (with GO_MF and MOP having the smallest amount of training data, see tables 1 and 2). To test whether the simplest model parameters can be used for more complex models (thereby saving a significant amount of resources), the same parameters were used for the BiLSTM-CRF and Char-Embeddings models, along with modified values for BiLSTM-ELMo due to resource constraints as discussed below.

The BiLSTM-ELMo model required the most resources for training and performed the worst. Due to memory issues using the simplest model parameters, we needed to restrict the batch size to 18 and thus took the best results from the tuning of the aforementioned BiLSTM with this batch size (see Table 14). Using a batch size of 18 (the smallest batch size we tested) resulted in some differences in the optimal numbers of epochs and neurons compared to the BiLSTM results shown in Table 13. Most ontologies still required 10 epochs for optimal performance, but a few (GO_CC, GO_MF and MOP) changed from 10 to 100 or vice versa, with the rest remaining the same. For neurons, most ontologies remained at 12, but two switched to 3, including ChEBI and GO_CC, with GO_MF remaining at 3 as well. Comparing F1 scores to the previous algorithms, the micro-F1 scores are exactly the same as those for the BiLSTM, but the macro-F1 scores either decreased slightly (for ChEBI, CL, GO_BP, GO_CC, GO_MF, and SO) or stayed the same (for MOP, NCBITaxon, PR and UBERON) as those for the BiLSTM. UBERON is the only ontology for which the parameters remained the same from the BiLSTM to BiLSTM-ELMo because it already required a batch size of 18 for optimal performance.

### Concept Normalization Results

Overall, exploring concept normalization as a translation problem through OpenNMT performed well on almost all ontologies in both the core and core+extensions sets (see Tables 15 and 17) and significantly outperforms ConceptMapper, except for PR and NCBITaxon_EXT, on the 30 held-out document evaluation set. Note that for ConceptMapper, not all text mentions were normalized to ontology class IDs, indicating false negatives, whereas for OpenNMT all text mentions were normalized to class IDs. These false negatives are the main reason performance of ConceptMapper is so poor (*e.g.*, no class IDs are predicted for MOP). Recall that ConceptMapper is a dictionary-based lookup tool where any text mention not in the dictionary will not be captured. Thus, methods like OpenNMT are necessary for unseen text mentions and perform quite well over previous dictionary-based methods.

For this task, evaluation can be performed at both the class ID level, in which a correctly predicted class ID in its entirety counts as an exact match, and at the character level, in which each character of the predicted class ID is evaluated, and a fractional score is calculated for a partial match. For example, while a text mention of “Brca2” was annotated in the gold standard with the Protein Ontology class PR:000004804, it was predicted to refer to PR:000004803, which differs from the gold standard only in the last digit. (The predicted ontology class ID PR:000004803 represents BRCA1, which also differs from the gold-standard protein BRCA2 in only its last digit.) At the class ID level, this is a mismatch, but at the character level it receives a score of 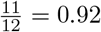, as 11 of the 12 predicted characters match. All ontologies performed above 69% on the class ID level, except for PR and PR_EXT. Comparing the core set to the corresponding core+extentions set, the extension sets outperform the core set for half of the ontologies (ChEBI_EXT, CL_EXT, GO_CC_EXT, PR_EXT, and UBERON_EXT). The class IDs of the extension classes may be easier to predict because there is less variation of text mentions annotated with the extension classes and also possibly due to the use of English words in the extension class IDs (*e.g.*, ChEBI_EXT:calcium, CL GO_EXT:cell) rather than the numeric IDs of the proper OBO classes. On the character level, all match scores are above 69% for both sets. The core set generally outperformed the corresponding core+extension set, except for (CL_EXT and GO_CC_EXT). For all ontologies, in most cases, the character level match scores are near or above the class ID ones, indicating that many of the predicted IDs were partly correct (in that they contained correct substrings). Future work can look at these partial matches for improvements.

To understand how well this method will generalize beyond the training data, we also report performance on the evaluation set with text mentions seen in the training data and text mentions unseen in the training data, again by full class ID and character level exact matching (see Tables 16 for the core set and 18 for the core+extensions set). Overall, for both the core and core+extensions sets on the class ID level, the text mentions seen in the training data performed far better than the unseen ones. Although, MOP, SO, MOP_EXT, and SO_EXT normalized correctly over 50% of the unseen text mentions. Only NCBITaxon, PR, and PR_EXT normalized none of the unseen text mentions. On the character level, the unseen text mentions results are still lower than the seen text mentions, but less of an extreme difference. Thus, these models can generalize to some extent beyond the text mentions seen in the training data, most likely due to the character-level translation of OpenNMT, unlike ConceptMapper.

To better understand these results and elicit future research directions, we looked at the types of mismatches that occurred at the class ID and character level. At the class ID level, two scenarios arise: (1) A different real class ID is predicted or a completely non-existent class ID is predicted. The aforementioned Protein Ontology example above, involving Brca1 and Brca2, is an example of the first scenario. This also occurs with concepts in the ontology hierarchy, such as the prediction for “monoatomic ion” (ChEBI:24867) to the ID for “monoatomic cation” (ChEBI:23906). One can see the striking resemblance between the two concepts; the only difference is the added “cat”, which OpenNMT did not pick up on. However, we do receive partial credit in the final evaluation as a monatomic cation is a monoatomic ion in the ChEBI ontology hierarchy. The second scenario happens much less frequently and includes examples, such as labeling text that should be annotated with SO class SO:0001179 (representing U3 snoRNA) instead with the class ID SO:0002000, which is not a real class ID. Note that these class IDs are quite different. The similarity between the predicted and the true class IDs are not always like Brca1 and Brca2, but can be wildly different class IDs. To understand how often both of these phenomena occur, we report the percentage of non-existent class IDs on the training, validation, and evaluation data out of the total mismatches (scenario two) for both the core and core+extensions sets (see Tables 19 and 20, respectively).

Overall, there is not much difference between the percentage of non-existent class IDs for the training and validation sets. However, there does seem to be quite a difference between those and the evaluation set, with more non-existent class IDs in the evaluation set (especially for GO_MF with 50%, and GO_CC_EXT and NCBITaxon_EXT with over 20%). At the same time, CL has no non-existent class IDs in any of the datasets and MOP_EXT has none in training and evaluation. This is a proxy for whether these ontologies can stay within the “vocab” of the class IDs (0% means fully within vocab). At the moment, it is unclear how to interpret these non-existent class IDs and we will explore this more in the discussion.

Focusing only on the core set, to determine the effect of concept frequency in our training data, we explored using the type-ids instead of the token-ids and it appears they have a modest affect on the performance of OpenNMT (see Tables 21 and 22 for class ID and character levels, respectively), except for NCBITaxon and PR. Interestingly, the token-ids compared to the type-ids generally stayed the same (CL, GO_MF, and SO), or slightly decreased for both the class ID and character level, except for MOP for class ID and GO_CC for character level where the type-ids performance increased. Note that on the class ID level, PR is very low (under 10%) to start for token-ids, so any drop is comparatively minuscule. For NCBITaxon, performance completely drops to zero on the class ID level, indicating that frequency of annotation concepts (token-ids) is necessary for performance (discussed more in the discussion section). However, on the character level, the drop is not as drastic for NCBITaxon due to partial matches. Looking at some of the mismatches for the type-ids compared to the token-ids, they do make different errors. For example, “penicillin” (ChEBI:173334) was predicted to be ChEBI:7986, which is the class ID for “pentoxifylline”. These class IDs are nothing alike, but we can see similar structure in the English concept mentions themselves with “pen” and “in,” that the token-ids identified, and the type-ids did not. This together suggests that the token-ids can be used over the type-ids for all ontologies except for MOP where using type-ids is better.

We also found that through our two other experiments (shuffled-ids and random-ids), the class IDs have some semantic content or structure (see Tables 21 and 22 for shuffled-ids and random-ids performance). As expected, we do see a decrease in performance from type-ids to shuffled-ids to random-ids for both the class ID and character levels for at least half of the ontologies. ChEBI, NCBITaxon, PR, and UBERON reduce to 0% for both shuffled-ids and random-ids on the class ID level, whereas CL, GO_CC, GO_MF, MOP, NCBITaxon, and SO generally maintain their level of exact match between type-ids, shuffled-id, and random-ids, with random-ids performing the best for CL of all experiments. Note that NCBITaxon is already at zero for the type-ids. Further, since OpenNMT works on the character level, we can see the structure breaking on the character level, where for all ontologies there is a large decrease from type-ids to shuffled-ids to random-ids, except GO_CC, GO_MF, MOP, NCBITaxon, and SO. This may suggest some accidental structure in the shuffled-ids and/or random-ids generated. Taking the class ID and character levels together, this suggests some structure in the current ontology class identifiers that OpenNMT has identified.

At the same time, we can add in more semantic content than the current ontology identifiers by alphabetizing the text mentions to see if it boosts performance for some ontologies (alphabetical-ids column in Tables 21 and 22). On the class ID level, the token-ids perform the best for all ontologies; however the alphabetical-ids recover the loss from the shuffled-ids and/or random-ids to perform close to the token-ids for most ontologies, except for CL, GO_MF, MOP, and NCBITaxon, with a similar trend on the character level. This suggests that the alphabetical-ids may provide too much structure that is imposed on the ontology concepts as we give an alphabetical-id to all text mentions even if they have the same class ID. For example, the class ID GO:0097617 pertains to the text mentions of annealing, hybridization, and hybridizations; and in the alphabetical-ids these were mapped to GO_MF:05728, GO_MF:15701, and GO_MF:15702, respectively. In the predictions then, hybridization was mapped to GO_MF:15702, which is one number different but incorrect nonetheless, whereas annealing was predicted correctly. Thus perhaps using stemming and lemmatization to provide one alphabetical-id to both hybridazation and hybridizations, while maintaining separate ids for annealing may boost performance.

## Discussion

By reframing concept recognition as a translation problem, we not only sidestep the multi-class classification problem, but also achieve above or near state-of-the-art performance (see Tables 5 and 6) on the concept annotation task of the CRAFT Shared Task with direct comparison to Furrer et al. [4] via the corresponding evaluation framework. Overall, on the full system run for the core set, our approach using BioBERT with OpenNMT slightly outperformed the best run from the participants in the CRAFT Shared Task (UZH@CRAFT-ST) [4] for six of the ten ontologies, whereas the latter modestly outperformed our best system for three of the four other ontologies (except for PR which was significantly lower). Further, the BioBERT used here required less resources than the BioBERT for UZH@CRAFT-ST, but the OpenNMT in our system made up for that (see Table 7). For the core+extensions set, our best-performing run slightly outperformed the best UZH@CRAFT-ST run for CL_EXT, GO_CC_EXT, and MOP_EXT, while the UZH@CRAFT-ST system outperformed our approach for all other ontologies. However, our BioBERT+OpenNMT system attained F1 scores within 0.2 of the UZH@CRAFT-ST scores for six of the seven ontologies with PR_EXT as the exception.

The errors that lead to lower performance in these full runs can come from multiple sources: span detection, concept normalization, and the interaction between them. For span detection, there are four classes of errors in relation to the gold standard text mentions: (1) the text mention is not detected at all; (2) the text mention is partially detected; (3) the text mention detected includes extra text; and (4) a full extra text mentions is detected. Case (1) is a false negative and case (4) is a false positive. For case (2) and (3) though, partial credit is awarded in the full evaluation pipeline for detecting at least part of the text mention. For concept normalization, the errors produced are most likely rather opaque and difficult to analyze (such as the non-existent class IDs) compared with *e.g.*, a dictionary-based approach like ConceptMapper (though there may be other reasons this approach would have trouble for these ontologies). Thus, concept normalization as translation needs to be explored beyond the experiments done here, especially since for PR and NCBITaxon_EXT, ConceptMapper outperformed OpenNMT. At the same time, some possible reasons for poorer performance may be related to the mixing of two different types of class IDs that include both numbers and English text as well as the varying lengths of the class IDs. The text class IDs tend to be rather long and OpenNMT performs worse with longer sequence lengths compared to shorter ones. However, the results on the core+extensions set, where many extension classes have English text class IDs and are longer, may provide some evidence that both the English text and longer class IDs perform worse than numeric and shorter class IDs as UZH@CRAFT-ST outperformed BioBERT+OpenNMT for all but three ontologies. Lastly, for the interaction of the two tasks in the full system, case (1) in span detection will lead to no normalization to a class ID, propagating the error from span detection and resulting in a false negative also for concept normalization. For case (2) and (3) however, it is possible for the normalization step to still correctly identify the class ID even though the text mention is slightly incorrect. On the other hand for case (4), the class ID determined will always be wrong because the text mention should not have been detected in the first place (OpenNMT always outputs a class ID for all text mentions inputted).

Overall, span detection algorithms performed very well for all ontologies with all F1 scores for the best algorithms above 0.81 for the core set and 0.77 for the core+extension set (see Tables 8 and 9). The lowest PR_EXT, seems to be suffering from predicting extra text mentions (false positives) as in case (4) of the span detection errors mentioned above. For concept normalization, the exact match percentage on the class ID level is above 72% for the core set (except PR) and above 69% for the core+extensions set (except PR_EXT) (see Tables 15 and 17). In this case, both PR and PR_EXT are very low at 10% and 15%, respectively, for class ID level. On the character level, the exact match percentage is much higher at 76% and 72%, but these are still lower than most of the other ontologies. Looking at the errors produced for these two ontologies specifically, it seems that the English text class IDs error mentioned above is at play for PR_EXT especially, as well as the fact that PR has longer numeric class IDs to begin with and it has many acronyms compared to other ontologies. As for the interaction of the two tasks, the best F1 scores of the full system were all above 0.69 for the core set and above 0.74 for the core+extensions set, except again for PR and PR_EXT, respectively (see Tables 5 and 6). It seems that the very poor results of concept normalization for PR and PR_EXT are to blame for these full system poor results, where many of the correct spans are detected but normalized incorrectly. Thus, there is still room to improve concept recognition for all core and extension ontologies, especially for PR and PR_EXT, and future work can directly compare to this work using the CRAFT Shared Task as their framework with the built in evaluation platform [51, 52].

Not only is state-of-the-art performance the goal, but also efficient use of resources. If the resources needed to train, tune, test, or use these models is outrageous then it would not be feasible to extend beyond the ontologies here or use these models for much larger collections of texts such as PubMed. Thus, some models are less useful than others due to their resource consumption for tuning and training (see Table 7), even if they perform better. Recent research quantified the exorbitant financial and environmental cost to deep learning algorithms for natural language processing [75]. They suggest that since researchers have unequal access to computational resources, researchers should first have equal access to computational resources, they should report their training time and hyperparameter sensitivity and perform a cost-benefit analysis (the benefit in terms of accuracy), and they should prioritize computationally efficient hardware and algorithms. For this work, having access to a GPU greatly sped up tuning and training times, which is in line with other research findings [74]. Compared to the BioBERT runs in UZH@CRAFT-ST [4], the CRF run on a CPU and BioBERT on a GPU were the most efficient algorithms because the CRF remains on the sentence level and BioBERT fine-tunes a pretrained language model (not taking into account the pre-training), respectively. If one does not have access to a GPU, or has a large dataset, the CRF would be most efficient for very similar performance, especially for GP BP, GO_MF, MOP, NCBITaxon, SO, GO_BP_EXT, MOP_EXT, and NCBITaxon_EXT, where the CRF either outperformed BioBERT or remained the same for span detection (see Tables 8 and 9). In terms of state-of-the-art language models, if one has a small dataset or access to a GPU, then BioBERT is preferred over the other language model, ELMo. ELMo requires the most resources and a GPU, and thus, may not be practical for this task, as the simple model parameters could not even be tested. This is in line with previous research comparing BioBERT and ELMo [45]. Along with ELMo, the BiLSTM may not be practical for this task as it requires a large amount of resources to tune (see Table 13). However a promising avenue for reducing BiLSTM resources is to reuse (or at least start from) the simple model parameters for more complex models with sometimes very slight drop or even an increase in performance. For concept normalization, OpenNMT also uses a large amount of resources (CPU threads, memory, and time), and other machine translation algorithms should be explored in the future with respect to resources and performance.

As mentioned above, for span detection, BioBERT performs the best, followed closely by the CRF for both the core and core+extensions sets. All algorithms that included a BiLSTM performed worse on this task. However at least one BiLSTM-included model performed well (within 0.2 of the best performing) for all ontologies except GO_CC and PR (see Table 8). There is always room to tune the BiLSTM parameters more, as it is difficult to tune them [63]. These results for BiLSTMs are in line with previous methods which found that BiLSTMs combined with other algorithms performed well on span detection [13, 30–32, 34, 35, 38, 40]. Thus, future work can explore the parameter reusage more, tune the algorithms more, and separetly tune BiLSTM-ELMo.

All algorithms, for at least one ontology, can also detect discontinuous concept mentions, which are one of the most difficult types of concept mentions to detect. A recent review on recognizing complex entity mentions, including discontinuous mentions, found that the problem is complex and not yet fully solved [16, 17]. We offer a new simple approach looking at the words between the discontinuous spans of the mentions, recognizing that they are greater in number compared to the words in the discontinuous spans. In the evaluation documents, we were able to detect some discontinuous spans for all ontologies even if only a few, except for ChEBI which contains the fewest discontinuous spans (see Tables 10 and 11). As reviewed by Dai [17], more system development as well as curation of more examples of these complex mentions is necessary to improve performance.

In general, regarding concept normalization as sequence translation performed well on all ontologies except for PR and PR_EXT as discussed earlier (see Tables 15 and 17). There are several advantages to this translation approach. The primary one is that the output to be predicted is a relatively short string of characters. This task is no longer a massive multi-classification problem with a choice among thousands of different classes. Treating the inputs as a sequence of characters, rather than a sequence of words also addresses the problem of unknown or out-of-vocabulary text mentions, as the model can learn sub-word patterns, covering potential text mentions in the evaluation set that are unseen in the training set, whose fragments (character n-grams) may appear in the training process [76] (see Table 2 for the number of new text mentions). However, poor performance in general most likely stems from the added ontology concepts from the OBOs not seen in the CRAFT annotations (see Table 1 for column additional OBO concepts), which greatly increased the number of concepts (especially for NCBITaxon and PR). However, performance for NCBITaxon is fine, whereas it is not for PR. For NCBITaxon this is the case because CRAFT mainly focuses on the laboratory mouse in biomedical studies. Thus, the mouse and human are overrepresented Taxa in this collection owing to the good results for token-ids (annotation frequency included) and the very bad results for type-ids (only one mapping). This suggests that for large ontologies using type-ids is not feasible, unless there is added training data for the most represented concepts in the corpus to boost the performance. For NCBITaxon specifically, this may mean that its model cannot generalize outside of the laboratory mouse biomedical studies articles. For PR on the other hand, there is no one overrepresented protein, like NCBITaxon, and so the large amount of additional OBO concepts is most likely confusing the model: it seems that the more annotation classes there are, the harder it is to train the model successfully. Note that ChEBI and GO_BP are the next largest ontologies and there performance does not suffer as much as PR and NCBITaxon for type-ids. Thus although, we added these extra ontology concepts for generalizability, a further exploration of the performance with different quantities and sets of additional ontology concepts not in CRAFT, such as concepts in other biomedical corpora, may help determine a better training dataset for concept normalization for some ontologies especially.

Combining span detection and concept normalization in the full run, our performance is near or slightly above the state of the art. One explanation is that the full evaluation pipeline gives partial credit to predicted IDs of ontology classes that have some semantic overlap with the correct annotation classes (*e.g.*, the aforementioned example involving CHEBI:‘monoatomic ion’ and CHEBI:‘monoatomic cation’). This may also be due to the duplicate concepts in the token-ids in the training data, as we included all annotations from CRAFT. However, it is interesting to note that the performances of the type-ids experiments are always the same or lower than that for the corresponding token-id experiments, except for MOP which is higher (see Table 21). The duplicates in the token-ids may bias the algorithm to recognize more frequent concepts in CRAFT. On the other hand, we do have a number of non-existent class IDs that are generated in almost all ontologies except CL, with more in the core+extensions set compared to the core set (see Tables 19 and 20). A further exploration of both these non-existent class IDs and the absence of them in CL may help explain how OpenNMT translates concept mentions to class IDs. Our results show that the unique ontology class identifiers contain some semantic content or structure (see Tables 21 and 22), which probably give rise to the non-existent class IDs in the full run. This is despite the recommended OBO Foundry identifier policy of identifiers not having semantic content [73]. There exists obvious semantic content in the IDs of a small number of proper OBO classes (*e.g.*, NCBITaxon:phylum, representing taxonomic phyla) and the many CRAFT extension classes of the OBOs (*e.g.*, CHEBI_EXT:calcium, representing elemental calcium and calcium ions) that contain English words. Semantic content may also arise due to a curation process that produces groups of closely related classes with sequentially adjacent identifiers, *e.g.*, PR:000004803 and PR:000004804, representing the BRCA1 and BRCA2 proteins, respectively. It appears that OpenNMT creates a fuzzy dictionary matching, where generally the mappings are unique with some wiggle room when necessary. By shuffling or randomly assigning class IDs, we can see where this fuzzy dictionary breaks the structure both on the class ID level as well as the character level. It appears OpenNMT is finding patterns between mentions of concepts with consecutively numbered class IDs, such as those for BRCA1 and BRCA2. In fact, many of the mistaken class IDs are only a few characters off from the correct ones (see Tables 15 and 17 on the character level). This therefore provides an opportunity for adding post-processing techniques on the class IDs to fix the mistaken characters, similar to the work of Boguslav et al. [25]. From another perspective, we attempted to boost performance by adding in more structure to the class IDs during the execution of the task, as in the alphabetical-ids experiment. The alphabetical-ids did not outperform the token-ids and the performance is most likely dependent on how alphabetical the concepts in each ontology are. For example, the concepts “HaC” (CL:0000855) and “haematopoietic cell” (CL:0000988) have different class IDs but are very close in the alphabet. Further, stemming and lemmatization may also help boost performance as mentioned previously. Thus, a hash or mapping from the current ontology class IDs to the alphabetical IDs with some stemming and lemmatization, along with some post-processing techniques on the character level, may help boost the performance on the class ID level. Both of these avenues warrant further investigation.

The main limitations of this work were due to limited data and resources. Beginning with the representations of the inputs for each task: for span detection the BIO-tags proposed here do not really capture overlapping spans nor capture many discontinuous spans. Future work on both of these is necessary. For concept normalization, we chose not to change any of the class IDs, even though they all had varying lengths within and between ontologies (particularly for the extension classes where the IDs included many more English text and were longer). Exploring different representations, including different ways to map the text class IDs to number identifiers, may improve performance. In terms of the algorithms themselves, future work should include an exploration of the amount of data needed to train and develop these algorithms for this task. In general it is possible to tune all algorithms more fully, including tuning the learning rate, where we kept the default or suggested for all algorithms. The BiLSTM experiments especially, were most likely not tuned fully and should be tuned additionally in future work, including using early stopping for determining a more precise number of epochs. Furthermore, the assumption that the two best-performing algorithms on the core annotations set for span detection would perform well on the core+extensions set may be false. Thus, future work should focus on the core+extensions set more for span detection, starting with the BiLSTMs. For concept normalization methods, we only explored one machine translation algorithms, OpenNMT, providing preliminary evidence that reframing this problem as translation is a salient avenue to explore in future work on concept recognition. Further exploration of other algorithms for machine translation may prove fruitful. Lastly, the unseen text mentions in the evaluation set hint at the generalizability of the concept normalization method, but the generalizability of both the span detection methods and the full end-to-end system to other biomedical corpora is unknown. It is unclear how to add synthetic data for span detection, but we did make sure to add all the class identifiers from the OBOs not seen in the training data to concept normalization. Future work should test the generalizability of both the span detection models and the full end-to-end system.

## Conclusions

In conclusion, machine translation is a promising avenue for concept recognition that sidesteps the traditional multi-class classification problem. We can achieve state-of-the-art results on the concept annotation task of the 2019 CRAFT Shared Task with a direct comparison to previous results. Given the amount of work that goes into shared tasks, shared task resources should be used and reused if possible. Further, resources in general need to be taken into consideration for concept recognition and NLP at large. Future work should focus on the core+extensions annotations set more for span detection. Also, for concept normalization, further exploration of other algorithms for machine translation may prove fruitful. As the generalizability of this system is unknown, future work should test the generalizability of the full end-to-end system.

## Declarations

### Ethics approval and consent to participate

N/A

### Consent for publication

N/A

### Availability of data and materials

The datasets used and analysed during the current study are available from https://sites.google.com/view/craft-shared-task-2019/home?authuser=0 (CRAFT Shared Task 2019).

CRAFT v3.1.3 was used and can be found here: https://github.com/UCDenver-ccp/CRAFT/releases/tag/v3.1.3.

The current CRAFT release can be found here: https://github.com/UCDenver-ccp/CRAFT.

All analyses and code are made available here: https://github.com/UCDenver-ccp/Concept-Recognition-as-Translation.

### Competing interests

The authors declare that they have no competing interests.

### Funding

This work is supported through NIH funding from three grants. NIH grant T15LM009451 support MRB and NDH. NIH grant R01LM008111 supports NDH, MB, WAB, and LEH. NIH grant OT2TR003422 supports MB, WAB, and LEH.

### Author’s contributions

MRB was responsible for the design and implementation of the project, including training and evaluating all models. NDH began this project as his thesis work determining the main ideas and framework for this project that were adapted by MRB. MB supervised all of the concept annotation work for the CRAFT Corpus as well as provided guidance and extensive discussions about the use of the corpus in this work. WAB created the CRAFT Shared Task with the evaluation pipeline, as well as provided guidance and extensive discussions on the shared task and computation for this work. LEH supervised the whole project.

## Acknowledgements

We would like to acknowledge the BioFrontiers Computing Core for computing resources and support, especially Jonathon Demasi. We would also like to acknowledge Harrison Pielke-Lombardo for preparing the CRAFT corpus annotations in a convenient format, as well as Asmelash Hadgu who supported Negacy Hailu and his thesis work that led to this manuscript. We acknowledge Lenz Furrer and Fabio Rinaldi for providing us information on their research. Also thank you to Katherine Sullivan for editing the manuscript.

## Notes

### Competing Interest Statement

The authors have declared no competing interest.

